# Targeting circadian transcriptional programs through a cis-regulatory mechanism in triple negative breast cancer

**DOI:** 10.1101/2024.04.26.590360

**Authors:** Yuanzhong Pan, Tsu-Pei Chiu, Lili Zhou, Priscilla Chan, Tia Tyrsett Kuo, Francesca Battaglin, Shivani Soni, Priya Jayachandran, Jingyi Jessica Li, Heinz-Josef Lenz, Shannon M. Mumenthaler, Remo Rohs, Evanthia Roussos Torres, Steve A. Kay

**Affiliations:** Alfred E. Mann Department of Biomedical Engineering, University of Southern California, Los Angeles, CA, USA; Department of Neurology, University of Southern California, Los Angeles, CA, USA; Department of Quantitative and Computational Biology, University of Southern California, Los Angeles, CA, USA; Division of Medical Oncology, Norris Comprehensive Cancer Center, University of Southern California, Los Angeles, CA, USA; Department of Statistics and Data Science, University of California Los Angeles, Los Angeles, CA, USA; Ellison Institute of Technology, Los Angeles, CA, USA

## Abstract

Circadian clock genes are emerging targets in many types of cancer, but their mechanistic contributions to tumor progression are still largely unknown. This makes it challenging to stratify patient populations and develop corresponding treatments. In this work, we show that in breast cancer, the disrupted expression of circadian genes has the potential to serve as biomarkers. We also show that the master circadian transcription factors (TFs) BMAL1 and CLOCK are required for the proliferation of metastatic mesenchymal stem-like (mMSL) triple-negative breast cancer (TNBC) cells. Using currently available small molecule modulators, we found that a stabilizer of cryptochrome 2 (CRY2), the direct repressor of BMAL1 and CLOCK transcriptional activity, synergizes with inhibitors of proteasome, which is required for BMAL1 and CLOCK function, to repress a transcriptional program comprising circadian cycling genes in mMSL TNBC cells. Omics analyses on drug-treated cells implied that this repression of transcription is mediated by the transcription factor binding sites (TFBSs) features in the cis-regulatory elements (CRE) of clock-controlled genes. Through a massive parallel reporter assay, we defined a set of CRE features that are potentially repressed by the specific drug combination. The identification of *cis*-element enrichment might serve as a new concept of defining and targeting tumor types through the modulation of *cis*-regulatory programs, and ultimately provide a new paradigm of therapy design for cancer types with unclear drivers like TNBC.

## Introduction

Circadian rhythms comprise the observed 24-hour cycle of physiological activities of living organisms. At the molecular level in mammals, these rhythms are formed by inter-connected negative feedback loops of gene transcription and translation.^1^ In its most simplified form, two transcription factors BMAL1 (Brain and Muscle ARNT-like 1, also known as ARNTL) and CLOCK (Circadian Locomotor Output Cycles Kaput) form a heterodimer and activate the transcription of a large set of genes, termed clock-controlled genes (CCGs), including repressors of themselves such as PERs, CRYs, and REV-ERBs. These repressors are translated and relocated back to the nucleus to repress the transcriptional activity (CRYs and PERs) or expression (REV-ERBs) of BMAL1 and CLOCK, thus forming the negative arm of the feedback loop. The alternating active and repressed cycles of BMAL1 and CLOCK activity form the master oscillator of circadian rhythms.^1^

Many essential physiological and cellular activities are under circadian control, such as the sleep-wake cycle and immune response on the organismal level, and metabolism, DNA repair, and cell cycle on the cellular level.^2,3^ Most human protein-coding genes are at least expressed in a circadian manner in at least one tissue.^4^ Therefore, it is not surprising that circadian genes are involved in hallmark pathways driving tumorigenesis.^5,6^

Although disruption of circadian rhythms on the organismal level by factors such as shiftwork has been a noted risk of cancer for decades, the molecular connections of circadian genes in cancer on the cellular level have only been established recently.^7,8^ In an early report of shRNA screening in acute myeloid leukemia (AML), BMAL1 and CLOCK emerged as oncogene hits, although detailed molecular mechanisms remain to be investigated.^9^ More recently, our laboratory and others reported that BMAL1 and CLOCK are indispensable in glioblastoma pathology. More specifically, BMAL1 gained a significant number of new chromatin binding sites in glioblastoma stem cells compared to non-tumor neural stem cells and altered the gene expression profile in multiple pathways including the TCA cycle.^10^ Application of small molecules that target BMAL1 and CLOCK in *in vitro* and *in vivo* models and showed their potential as experimental therapeutics for glioblastoma.^10^

These studies, among others, established that circadian clock proteins are actionable targets in cancer, demanding more detailed studies on their biology and clinical relevance. It is becoming clear that their functions can be altered in different ways and thus be oncogenic or tumor-suppressing in different tumor types.^6^ Therefore, to make most use of available drugs targeting circadian genes, more studies need to be done for defining a Target Product Profile including patient populations and designing therapeutic strategies.

In this study, we investigated the targetability of circadian gene-regulated transcription in breast cancer. Clinically, breast cancers are treated according to their hormone receptor (HR) and human epidermal growth factor receptor 2 (HER2) expression status, which are major drivers of the progression of corresponding cancer types.^14^ Those lacking both HR types (estrogen and progesterone) and HER2 are termed triple-negative breast cancer. TNBC is the most aggressive subtype of breast cancer, for which no recurrent drivers or therapeutic targets have been recognized currently, making it the subtype with the poorest prognosis. ^14^ It has been reported that the circadian rhythm in TNBC is disrupted, indicating altered circadian gene functions. We also focused our functional study of circadian genes on TNBC.

Our results showed that the positive arm genes, BMAL1 and CLOCK, in the circadian gene network are potential targets for mMSL TNBC cells. Using small molecule modulators, we repressed BMAL1- and CLOCK-regulated transcriptional programs to repress the proliferation of mMSL TNBC cells. Using high-throughput screening experiments, we revealed that BMAL1 and CLOCK might function through certain types of *cis*-regulatory elements (CREs) to drive the proliferation of mMSL TNBC cells. Repressing the activities of such CREs with small-molecule drugs might provide a new model for defining oncogenic drivers and therapeutic targets in breast cancer.

## Results

### Features of circadian gene expression in breast cancer clinical samples

To assess the clinical relevance of core circadian genes in breast cancer, we first analyzed their expression signatures in the TCGA and METABRIC cohort data.^11^

The major challenge to study circadian gene dynamics in clinical samples is that usually only a single time point of data is available for each patient, and the sampling time is mostly unmarked. Despite recent successes in reconstructing rhythmicity from unmarked patient data using neural networks^12,13^, constructing explainable biomarker signatures is still a challenging task. However, it has been shown that circadian rhythm is often disrupted or lost in cancer cells, as indicated by the loss of negative correlation between positive and negative arm genes in tumor tissues (Supplementary Fig. 1a and Li et al.^12^). This implies that at the population level, circadian rhythm is disrupted or at least diminished in breast cancer. We therefore examined their expression signatures in these datasets without factoring in time of sampling.

Firstly, by contrasting the normal and tumor tissue expression data in the TCGA cohort, we found that most differences in gene expression were in the negative arm genes (6/7, Supplementary Fig. 1b). They were downregulated in tumor tissues, whereas the positive arm genes showed either small or insignificant differences. By visualizing tumor versus normal tissues within the same patient group, clear trends of downregulation in tumors were observed for PER1, PER2, and CRY2 (Figure 1a). This result implies that there is a global pattern of change in the expression of core clock genes in breast cancer. To identify a “tumor signature” of clock genes, we ran logistic regression on the expression data from the cohort with paired tumor and normal tissue (n=114) and then tested this model on the remaining tumor-only samples (n=982, Figure 1b). The model resulted in a 6% training error and an 11% testing error, and the coefficients of significant factors (p < 0.01) generally agree with the differential expression gene trends. These results confirmed that in breast cancer, circadian clock genes have a traceable pattern of alteration in tumors, and the negative arm genes are repressed at large.

**Figure 1.**
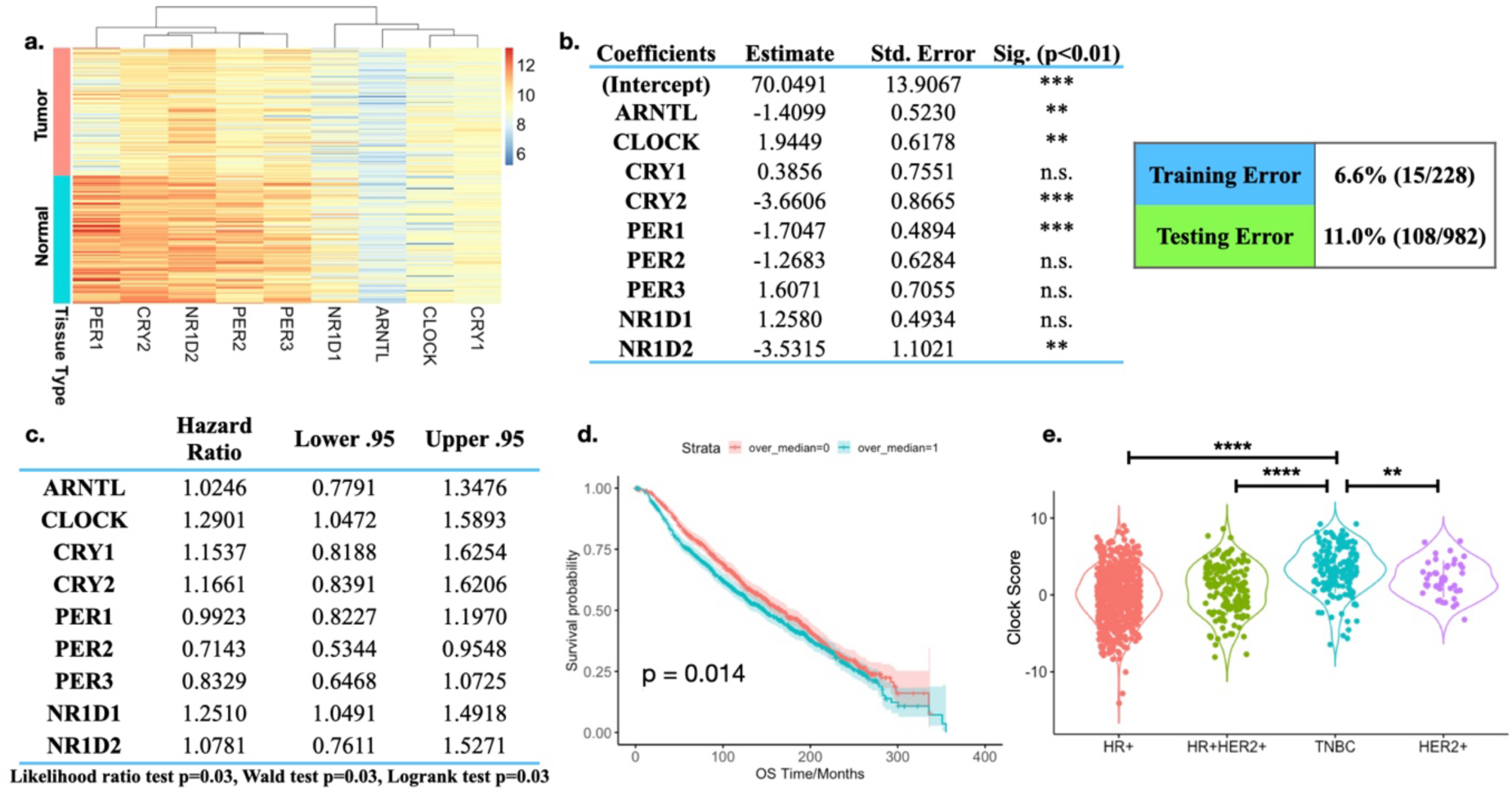
Clinical significance of circadian clock genes in breast cancer. **a.** Parallel comparison of core circadian gene expression in normal (green) and tumor (pink) tissues in the TCGA cohort. Only patients with both tumor and normal tissue available are plotted. **b.** Results of logistic regression model trained on paired tumor-normal patient samples to distinguish tumor tissues from normal tissues. **c.** Results of multivariate Cox-regression model trained on core circadian genes. **d.** Validation of TCGA-trained Cox hazard ratio model on METABRIC cohort data. p-value was calculated by log-rank test. **e.** “Clock score” of TCGA cohort data on different breast cancer subtypes. (*: p<0.05; **: p<0.01; ***: p<0.001; ****: p<0.0001.)

We then sought to test if clock genes have prognostic implications for breast cancer patients. For single circadian genes, when we stratified cohorts by their median expression level, only CRY1 showed a significant difference in overall survival across all samples (Supplementary Fig. 1c). Within subtypes, only REV-ERBα and PER3 in TNBC are significant (Supplementary Fig. 1d). To test if the “tumor signature of clock genes” from the logistic model also have prognostic implications, we calculated a “tumor clock score” by summing up the clock gene expression levels weighted by the logistic model coefficients, then tested it on both TCGA and METABRIC datasets. Interestingly, using the median score to stratify cohorts, this score did not generate a clear difference in survival (Supplementary Fig. 1e). To find a prognostic signature of clock genes, we ran multivariate Cox regression on the TCGA dataset. The Cox model successfully generated a risk signature (Figure 1c and Supplementary Fig. 1f), prompting us to test it in the METABRIC dataset as a validation. We summed clock gene expression levels weighted by coefficients from the Cox model and found that the higher-score group indeed had significantly shorter overall survival (Figure 1d).

Comparing the logistic and the Cox models, we found that high CLOCK expression is both a feature of tumors and a risk factor for overall survival. By contrast, the negative arm genes are suppressed in tumors. Together, these data revealed the clinical relevance of circadian genes and the potential of the positive-arm genes to support tumor progression in breast cancer.

The results from clinical data prompted us to consider how to extract a therapeutic hypothesis for leveraging clock protein modulation in breast cancer. We therefore tested whether the repression of the positive-arm clock genes will impact the proliferation of breast cancer cells.

To indicate the overall balance between positive- and negative-arm genes in the core circadian circuitry, we constructed a clock scoring system that sums the z-scores of the expression of positive arm genes (BMAL1 and CLOCK) and subtracts the z-score of negative arm genes (CRY1/2, PER1/2/3, and REV-ERBα/β). Among different subtypes in the TCGA samples, TNBC has the highest clock score (Figure 1e), implying highest activity of positive-arm genes. Because TNBC lacks known targets and available therapeutic strategies, there’s a more urgent need to find potential new targets. We therefore chose TNBC to test if positive-arm clock gene BMAL1 and CLOCK are potential therapeutic targets.

### Knockdown of BMAL1 and CLOCK disrupts proliferation of mMSL TNBC cells

TNBC is a highly heterogeneous disease containing various molecular subtypes defined by their molecular signature.^15^ To further test the dependency of TNBC cells on BMAL1 and CLOCK, we performed a shRNA screening on a panel of cells including TNBC cells across different molecular subtypes (Figure 2a) and two non-cancer cell lines (immortalized human mammary gland epithelial cell MCF10A and human lung fibroblast cell IMR-90, Supplementary Figure 2). From the initial screening, only mMSL TNBC cells showed visibly hampered proliferation (Supplementary Figure 2). We further quantified tumor cell proliferation with CellTiter-Glo. Following KD, mMSL cells showed significantly reduced proliferation after 4 days (4/4 shRNAs for MDA-MB-231 and MDA-MB-157, 3/4 shRNAs for MDA-MB-436), whereas other types either exhibited less significant effects or promoted proliferation (Figure 2b and Supplementary Figure 3a). This result is consistent with previous reports that BMAL1 and CLOCK are essential in stem-like cancer cells in glioblastoma and AML.^9,10^

**Figure 2.**
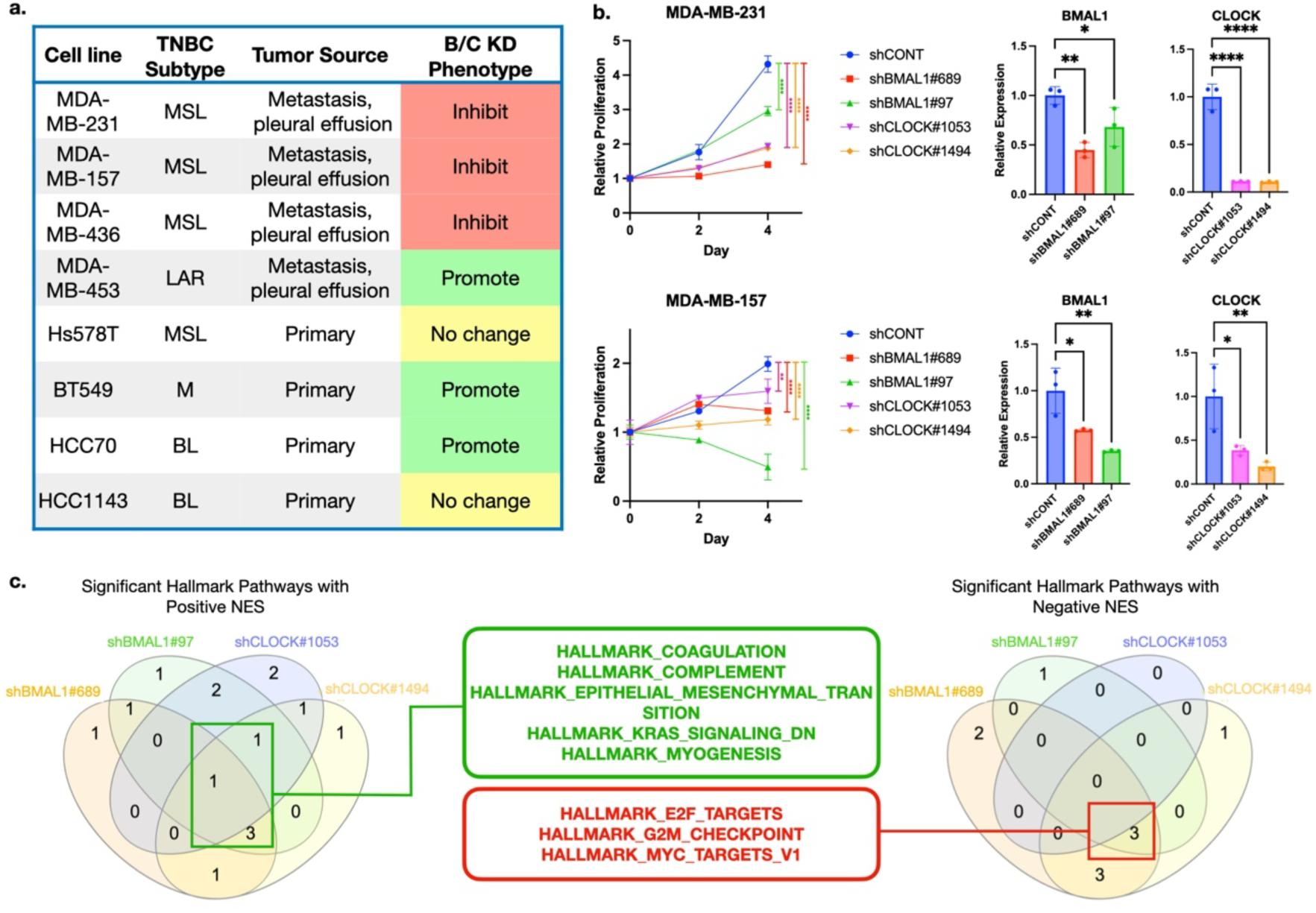
Knockdown of BMAL1 and CLOCK impairs the proliferation of metastatic mesenchymal stem-like TNBC cells. **a.** Summary of phenotype screening on a panel of TNBC cell lines crossing different subtypes. MSL: Mesenchymal Stem-like; LAR: Luminal androgen receptor; M: Mesenchymal; BL: Basal-like. **b.** Quantification of cell proliferation after BMAL1 and CLOCK knockdown using CellTiter Glo (Left Panel) and validation of knockdown efficiency using quantitative RT-PCR (Right Panel). p-value was calculated using one-way ANOVA. (n=4 for CellTiter Glo and n=3 for quantitative RT-PCR. *: p<0.05; **: p<0.01; ***: p<0.001; ****: p<0.0001.) **c.** Venn diagram summarizing GSEA results on Hallmark pathways from BMAL1- and CLOCK-knockdown MDA-MB-231 cells.

To understand the mechanism of repressed proliferation by the knockdown of BMAL1 and CLOCK, we performed RNA sequencing on BMAL1- and CLOCK-knockdown MDA-MB-231 cells (Supplementary Figure 3b). We ran GSEA analysis and compared the change in Hallmark pathways after KD (Figure 2c). We consider pathways that are enriched in at least three out of four shRNA groups the most relevant ones to BMAL1 and CLOCK KD. Interestingly, epithelial-mesenchymal transition (EMT) is the only pathway that is enriched in all four KD groups. Because only mMSL cells are sensitive to BMAL1 and CLOCK KD, this result further suggests that the EMT and the related stem cell feature are potential indicators of BMAL1 and CLOCK functioning as oncogenic genes and potential targets for therapy. On the other hand, all the enrichments with negative NES are cell cycle-related (E2F targets, G2-M check point, and MYC target genes). We validated this result by performing EdU assay and found reduced EdU-positive cell proportion following KD of BMAL1 (Supplementary Figure 3c). Because circadian rhythm and cell cycle are well coordinated process and B/C are important regulators of cell cycle, this result suggests that BMAL1 and CLOCK may contribute to cell cycle directly in the transcription level, although more detailed mechanistic confirmation would be needed.

### CRY stabilizer and proteasome inhibitors synergize to repress BMAL1 and CLOCK activity and proliferation in mMSL TNBC cells

To harness the tumor-supporting function of BMAL1 and CLOCK as potential therapeutic targets, we proceed to investigate if their activity can be targeted by small molecules in TNBC. Currently there is a lack of direct small-molecule modulators for BMAL1 and CLOCK, we thus used several small molecules that strengthen the negative arms of the clock network and in turn repress BMAL1 and CLOCK function. (Supplementary Figure 4a)

To test the small molecules’ effects on repressing BMAL1/CLOCK (B/C) activity and cell proliferation, we first compared their IC50 and their effect on the expression of canonical B/C target genes (DBP, CRY1, and PER1, Figure3a-b). REV-ERB agonist SR29065^16^ and CK2 inhibitor GO289^17^ have relatively low IC50, but the expression of B/C target genes was not suppressed. This indicates that in these TNBC cell lines that we tested, their viability effects are more likely due to mechanisms other than modulating B/C activity. On the other hand, CRY2 stabilizer SHP1705^18,19^ can faithfully repress the expression of DBP, CRY1, and PER1, but has less of an effect on cell proliferation than the other small molecules. Because our interest is in targeting the activities of B/C, we sought to understand and expand the function of SHP1705 via manipulation of the transcriptional potential of the BMAL1::CLOCK complex.

To confirm the effect of SHP1705 on B/C, we performed bulk RNA-seq on MDA-MB-231 cells treated with SHP1705 (10 µM) for either 8 or 24 hours. SHP1705 exhibited a relatively small effect on the mRNA transcription landscape resulting in a total of 1451 differentially expressed (DE) genes, of which only a handful had more than a 1.5-fold change (Figure 3c). However, it was highly selective towards genes in the core circadian regulatory network and repressed the negative-arm genes for at least 24 hours. (Figure 3c and Supplementary Figure 4b). Gene set enrichment analysis (GSEA) and gene ontology (GO) analysis confirmed that SHP1705 repressed core circadian gene network and had minor effects on hallmark pathways in TNBC, reflecting its target specificity (Figure 3d-e, Supplementary Fig. 4c). This focused selectivity of SHP1705 on core circadian gene network may not be sufficient to inhibit the transcription of other genes mediated, but not primarily determined, by B/C in mMSL TNBC cells. This may explain the limited effects that SHP1705 has on cell proliferation as a single agent.

**Figure 3.**
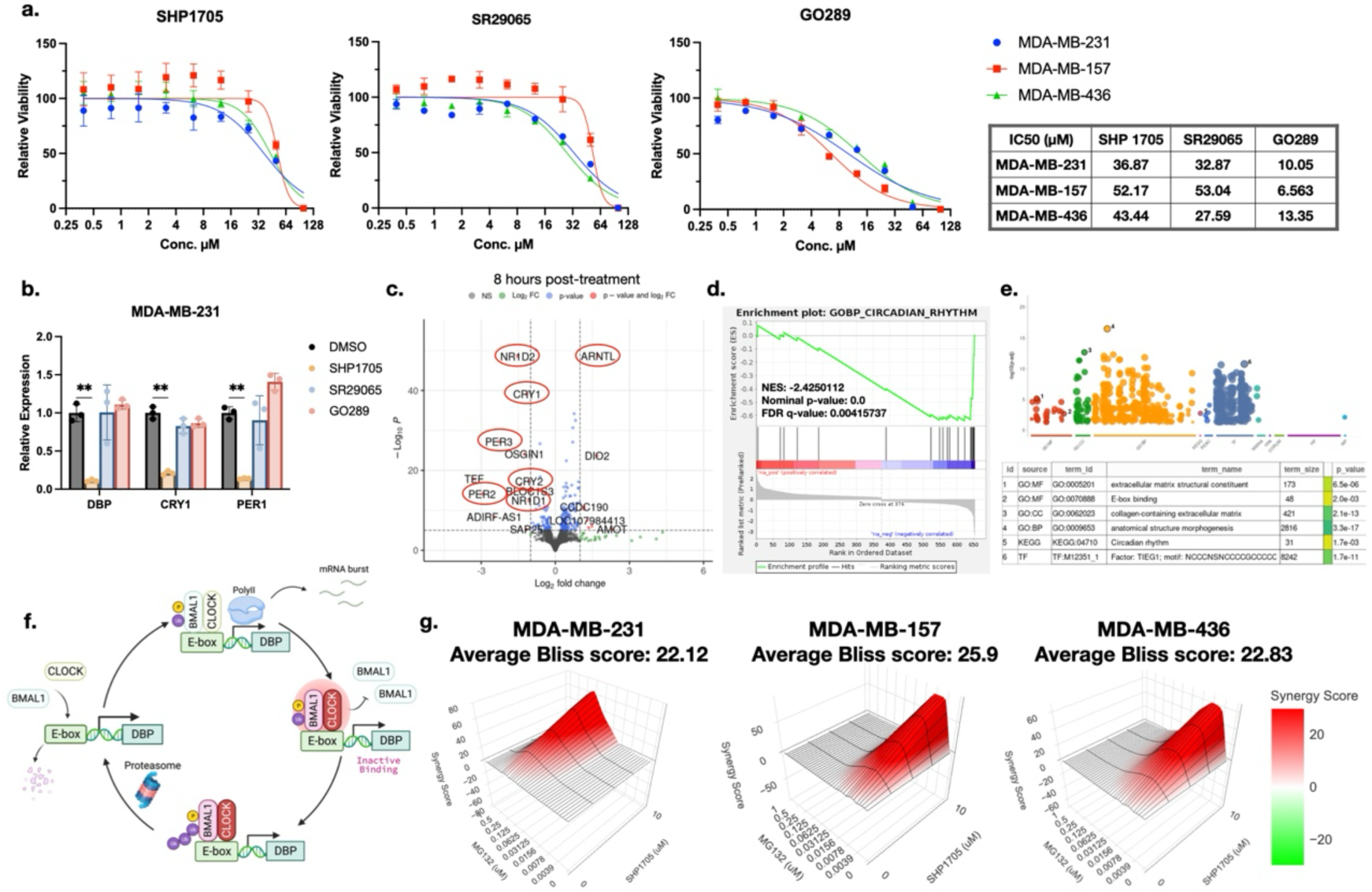
CRY stabilizer SHP1705 represses core BMAL1 and CLOCK target genes and synergizes with proteasome inhibitors. **a.** Response curve of small molecules that target different parts of the circadian gene regulatory network on the proliferation of mMSL TNBC cell lines. SHP1705 stabilizes CRY, SR29065 agonizes REV-ERBs, and GO289 inhibits CK2, three repeats were done for each data point. IC50 values of each small molecule in each cell line were summarized in the table. **b.** Gene expression level change of canonical BMAL1 and CLOCK target genes after 8 hours of small-molecule treatment (10µM SHP1705, 1µM SR29065, and 1µM GO289) quantified by quantitative RT-PCR, p-values were calculated using t-test, n=3. **c.** Volcano plot of differentially expressed genes in MDA-MB-231 cells after 8 hours of 10 µM SHP1705 treatment, core circadian clock genes are marked in red circles. **d.** GSEA analysis showing enriched circadian rhythm gene set after SHP1705 treatment. **e.** GO enrichment analysis showing enriched E-box-binding and circadian rhythm terms after SHP1705 treatment. **f.** Reported mechanism of proteasome function in BMAL1 and CLOCK transcription activity. In each cycle of transcription burst, inactive BMAL1 and CLOCK need to be removed by the proteasome from DNA in order for a new round of active BMAL1 and CLOCK binding to initiate another round of transcription burst. **g.** Synergy curve and average Bliss synergy score of SHP1705 and proteasome inhibitor MG132 on repressing three mMSL TNBC cell proliferation.

To improve upon the effects observed with SHP1705 treatment, we sought for other complementary modulators of B/C transcriptional activity. It was previously reported that the proteasome is a direct regulator of B/C transcriptional activity.^20^ In this model, once B/C initiates a round of transcription burst, the dimer becomes inactive on the DNA strand, preventing new rounds of active dimer from binding. Proteasomal degradation of B/C is necessary for new rounds of transcriptional bursting. (Figure 3f) We reasoned that if CRY2 and the proteasome both regulate B/C-mediated gene transcription programs, their effects would be synergistic. We tested two proteasome inhibitors (PIs), MG132 and carfilzomib (CFZ), with SHP1705 on all three mMSL cell lines. MG132 showed strongly synergistic effect (Bliss synergy score > 20) with SHP1705 in inhibiting cell proliferation (Figure 3g). CFZ also exhibit synergistic effect on two out of three cell lines (Supplementary Figure 4d).

In summary, we found that in mMSL TNBC cells, CRY2 stabilizer SHP1705 can selectively suppress B/C activity in regulating core circadian network genes and synergize with proteasome inhibitors to trigger reduced cell proliferation.

### Combination of SHP1705 and MG132 inhibits the circadian transcription program

To understand and leverage the mechanism of this synergy, we performed RNA-seq on single- and dual-drug-treated cells after 8 hours of treatment. We used 100 nM of MG132 and 10 µM of SHP1705, the highest concentrations of single drugs that have no significant effect on cell proliferation by themselves, but has strong effect combined. In the sequencing data, the expression levels of core BMAL1 and CLOCK target genes CRY1, PER1, and DBP after SHP1705 treatment are consistent with quantitative PCR data from previous section (Supplementary Figure 5a). This result provided a validation of the sequencing data. Gene set enrichment analysis on the drug combination data showed negatively enriched E2F targets and the G2M checkpoint (Figure 4a), which is consistent with the results in KD analysis (Figure 2c) and supports that the function of BMAL1 and CLOCK are repressed by the combination of MG132 and SHP1705.

**Figure 4.**
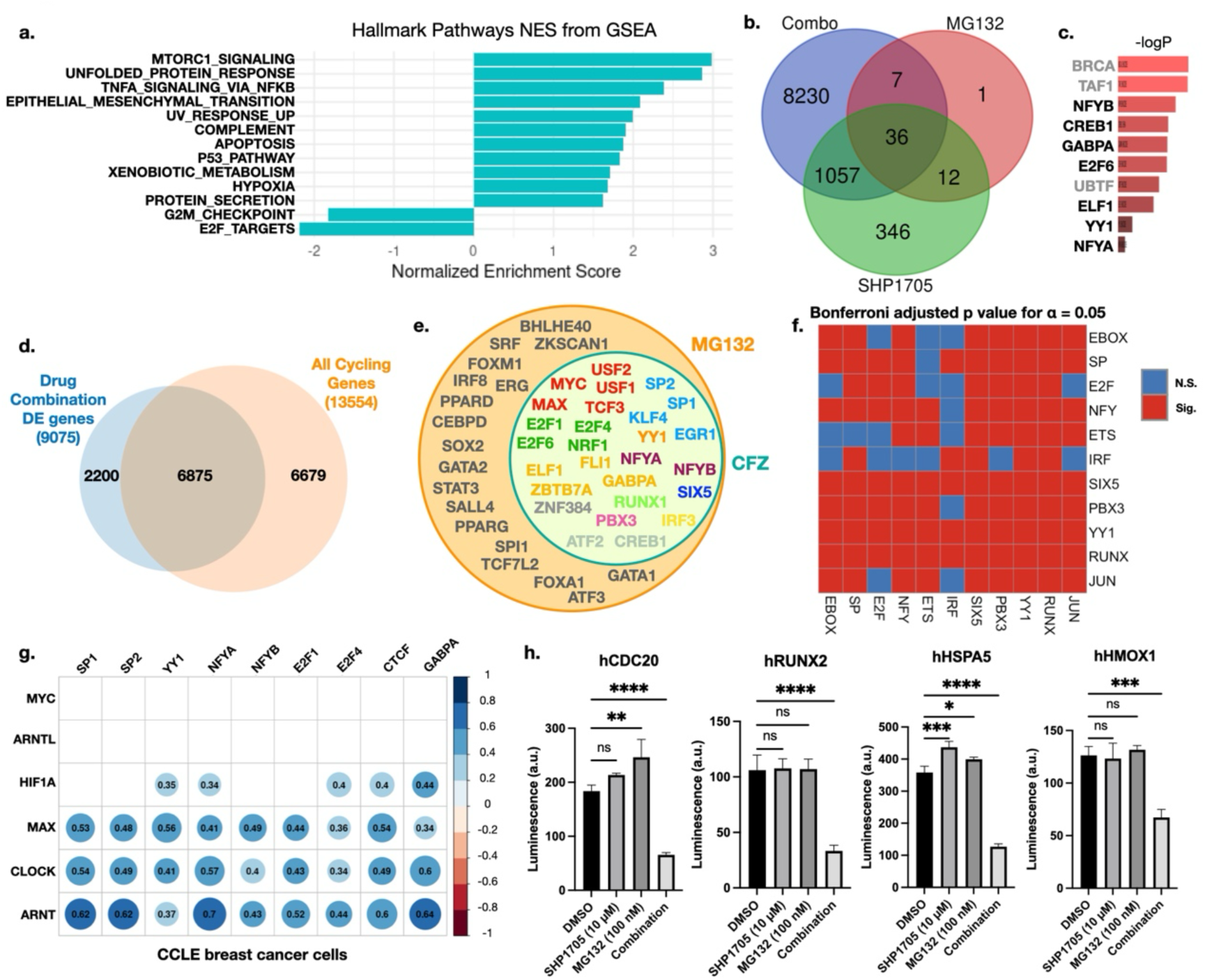
SHP1705 and MG132 combination repress the circadian transcription program through a cis-regulatory mechanism. **a.** Venn diagram shows the intersection of DE genes in single and dual drug-treated MDA-MB-231 cells. **b.** GSEA analysis on hallmark pathways shows negatively enriched G2M checkpoint and E2F target gene sets. Only significant results (FDR q-value <0.05) are plotted. **c.** Transcription factor (TF) targets enrichment on combination-specific DE genes shows that in addition to E-box-binding TFs, other enriched factors include reported CCG regulators such as ETS, YY, and NFY family factors. TF names showed in gray means they are either not *bona fide* TFs that binds DNA (BRCA and TAF1) or primarily regulates the transcription of non-mRNAs (UBTF). **d.** Venn diagram showing the overlap between DE genes in the drug combination group and reported circadian cycling genes. **e.** Intersection of enriched TFs from combination-specific DE genes in the MG132-SHP1705 and the Carfilzomib-SHP1705 group. The TFs that are shared by both groups are colored based on their binding motifs. **f.** Co-occurrence analysis of refined motifs in the H3K27ac peaks of MDA-MB-231 cells shows that the motifs indeed show a significant feature of cooccurrence in the active chromatin regions of MDA-MB-231 cells. E-box co-occurs with most other motifs found in Venn diagram analysis. **g.** Correlation analysis performed on all breast cancer cell lines in the CCLE database showed that the β-subunits of E-box-binding TFs and EBTF-Co factors show strong positive correlations. Size of the circles is correlated with the negative-log transformation of corresponding p-values, and the number in the circles represents Pearson correlation coefficients. **h.** Luciferase reporter assay for promoter activity showing that CRE activity of DE gene promoters in the combination group is repressed. (n=3, *: p<0.05; **: p<0.01; ***: p<0.001; ****: p<0.0001.)

Venn diagrams of DE genes in different treatment groups showed that at the dosage levels we used, SHP1705 and MG132 show little effect on the transcriptome of MDA-MB-231 cells, but their combination resulted in over 9,000 DE genes (Figure 4b and Supplementary Figure 5b-c). We thus assume that the synergistic effect is mediated by the DE genes that appear exclusively in the combination group but not in single drug-treated groups.

Given that this drug combination directly targets part of the transcription mechanism, we performed enrichment analysis on TF target genes to investigate the molecular mechanism (Figure 4c). To focus on genes that account for the synergistic effect, we ran an enrichment analysis on DE genes that appear exclusively in the combination group but not in single drug-treated groups. E-box-binding factors are still significantly enriched in the combination group, as in SHP1705-treated cells (Supplementary Fig. 5e), suggesting the role of E-boxes in this effect. The next most enriched TFs include NFY factors, YY1, ETS family, and E2F family factors (Figure 4c). Interestingly, the binding sites of these TFs are found overrepresented in the promoters of CCGs, and they thus are thought as potential combinatorial regulators of the circadian rhythmicity of gene transcription.^22^

This result implies that the drug combination expanded the repression of core B/C target genes by SHP1705 alone to a larger set of circadian-controlled genes. Indeed, by intersecting DE genes in the combination group and an annotated circadian cycling gene set,^4^ we found that most (6875/9075, 76%) of the DE genes are cycling genes, as predicted by our hypothesis (Figure 4d).

### SHP1705 and MG132 repress gene transcription through a cis-regulatory mechanism

To provide stronger evidence on our hypothesis, we conducted RNA-seq and performed the same analysis for the CFZ-SHP1705 combination (Supplementary Fig. 5e-f) and intersected the TF enrichment results from these two groups (Figure 4e). The refined TF list confirmed the potential involvement of E-box-binding factors (MYC, MAX, and USF1/2) and the TFs whose binding motifs are enriched in CCG promoters. Interestingly, we noticed that many of these TFs bind to the same DNA motif, allowing us to further refine the list of TFs to a set of binding motifs (Supplementary Figure 5g). Binding motifs of TFs are combined in CREs to determine their functions, this observation thus points to a possibility that by using the drug combination we repressed the activity of certain CREs whose functions are mediated by these *trans*-acting factors, including BMAL1 and CLOCK. Supporting this hypothesis, the list of motifs we found in this study largely overlaps with another list of motifs that are reported to comprise TNBC-cell-specific super-enhancers.^23^ These results suggest that these motifs may constitute a specific subtype of CREs.

To confirm that these motifs indeed are located near each other in the active CREs of TNBC, we analyzed Histone 3 Lysine 27 acetylation (H3K27ac) ChIP-seq data of MDA-MB-231 cells. By searching for clusters of these motifs, we found that these motifs indeed collocate in CREs and can show specific arranging features such as alternating SP1 motif and E-boxes in tandem. We also quantified significant co-occurrence of pairs of the motifs in the clusters and found that E-boxes tend to co-occur with most other motifs (Bonferroni adjusted p for significant level 0.05, Figure 4f).

Since these motifs are collocated in the CREs, we reasoned that if they function collaboratively, their binding TFs would have correlated expression levels as well. We therefore tested the correlation of E-box-binding factors (EBTFs) and TFs that bind to the hypothetical collaborative motifs, which we term hypothetical EBTF-Cofactors (EBTF-Co) for simplicity. In the TCGA-BRCA dataset, CLOCK and ARNT showed a moderate correlation with EBTF-Cos (Supplementary Fig. 5h). However, it is noteworthy that the statistical power may be reduced because of the involvement of non-tumor tissues such as stromal and immune cells in patient tumor samples. To alleviate the concern and focus on cancer cells, we performed the same analysis in datasets of breast cancer cell lines from the Cancer Cell Line Encyclopedia (CCLE).^24^ Interestingly, MAX, CLOCK, and ARNT showed strong positive correlations with all EBTF-Co’s, whereas MYC, BMAL1, and HIF1A did not (Figure 4g). Among these heterodimer pairs of b-HLH TFs, MAX, CLOCK, and ARNT are β-subunits whose expression level tends to be more stable and constitutive, whereas MYC, BMAL1, and HIF1A are α-subunits that are more dynamically expressed.^25^ Specifically, BMAL1 mRNA level is under strong circadian control, and MYC mRNA degrades quickly. This dynamic nature of the α-subunits might compromise their statistical power. Therefore, the correlation between EBTF β-units and EBTF-cofactors supports our hypothesis that they collaboratively contribute to the CREs that contain their binding motifs.

To test this hypothesis that the activity of CREs was repressed by the drug combination, we cloned the promoters of representative DE genes that contains the binding motifs of the EBTF-cofactors (Supplementary Fig. 6), and their activity was repressed after combination drug treatment as measured by a luciferase reporter assay (Figure 4h). These results confirmed that the drug combination repressed gene expression by inhibiting the activity of their promoters that contain the motifs of EBTF-Cofactors.

### The combination of SHP1705 and MG132 suppresses a circadian transcription program by inhibiting selected types of CREs

To define the features of the CREs that are repressed by the drug combination in an unbiased manner, we performed Self-Transcribing Active Regulatory Region Sequencing (STARR-seq) to test CRE activities in cells treated with single drugs or their combination (Figure 5a).^26^

**Figure 5.**
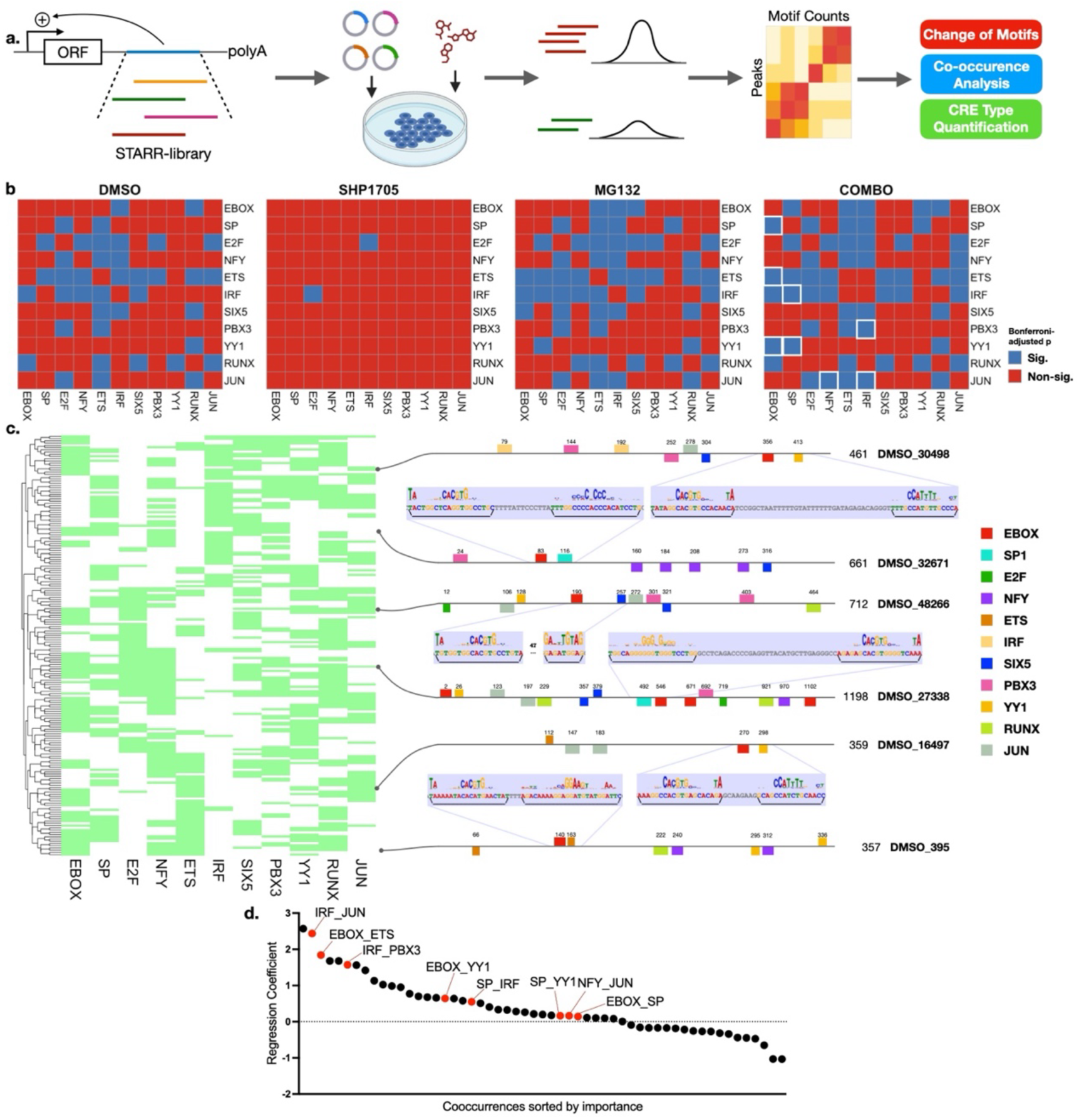
Massive parallel reporter assay reveals types of CREs that are potentially repressed by the drug combination. **a.** STARR-seq experiment design. Genomic library from MDA-MB-231 cell genomic DNA is inserted into the hSTARR-seq vector, and their CRE activity are quantitatively inferred as read abundance. Peaks are called for each treatment group and motifs in each peak are counted to generate peak-motif count matrices. Then motif analyses are implemented on the matrices. **b.** Motif co-occurrence matrices showing distinct features in different treatments. Comparing the combination (COMBO) and control (DMSO) groups, co-occurrences that were lost in the combination group are highlighted by a white frame. **c.** Summarized CRE “types” that are potentially repressed by the drug combination and detailed motif distributions in CREs from selective types. Each row in the heatmap represents a “type” of CRE defined by the existence of indicated motifs in it. Representative CREs belonging to the repressed types are plotted on the side, numbers on each motif represent their location in the CRE. Direction of the motif-representing color square represent the strand on which the motif is located. **d.** Contribution of motif pairs to being a “combination-repressed type”. Pairs that lost significance in co-occurrence analysis are colored in red.

We used fragmented genomic DNA of MDA-MB-231 cells as an input library (700 base pair-long on average, Supplementary Fig. 7a). We achieved a complexity of more than 10 million unique reads for all groups (Supplementary Fig. 7b), which theoretically can cover twice of the human genome (around 3.1 billion bp in length). The input copy number of each fragment cannot be accurately defined in this experimental setting, therefore a precise fold-change analysis of STARR activity across different treatments cannot be reliably performed. Instead, we tried to leverage the size of the library and compare the overall representation of motifs. We selected the most highly enriched fragments by peak calling and performed a qualitative comparison to define CRE types that are repressed by the treatment of drug combination.

We first examined the global TFBS landscape of all STARR-seq peaks across different treatments (Supplementary Fig. 7c). A total of 258 clustered non-redundant motifs (columns) were counted through each peak (rows).^27^ The result shows that even with a relatively loose stringency (80% of maximum match score for each position weight matrix), motifs are relatively sparse in the peaks, and the drugs did not cause a visible change in the global landscape of motifs in STARR peaks. The following analyses thus focus on the EBTF-Co motifs.

Firstly, we tested if the drugs changed the representation of each single motif by comparing their abundance which is normalized to library complexity and peak number in each treatment group. Indeed, the E-box and most EBTF-Cos were less represented in the combination group compared to the DMSO group (Supplementary Fig. 7d). This result suggests that the overall activity of E-boxes and the other motifs are potentially repressed by the drug combination. Interestingly, SHP1705 led to the over-representation of several motifs, but the mechanism is unknown and can be explored in future studies.

Next, we tested the pair-wise co-occurrence of each motif in different treatment groups. We first tested the co-occurrence of all 258 clustered motifs. In general, motifs tend to co-occur with only a subset of partners and show traceable patterns in a heatmap, although we observe an inflation of co-occurrence in SHP1705 group (Supplementary Figure 8a). This result supports that we can subtyping CREs based on their motif composition and test their functionality. Comparing the combination to the DMSO group, we found several significant co-occurrences that were lost, including E-box with SP, ETS, and YY1; SP with IRF and YY1; NFY with JUN; IRF with PBX3 and JUN (Figure 5b). These losses of significance imply that the enhancer activities of peaks containing these motifs together are under-represented in the combination group, thus constituting promoters that are likely to be repressed by the drug combination.

To catch more specific features of CREs that contain these motifs, we quantified the numbers of motifs in STARR-seq peaks from each treatment group and performed dimension reduction using principal component analysis (PCA, Supplementary Fig.8b). However, each PC accounts for the same variance as single variables and no clear cluster can be defined (Supplementary Fig 8c). This might be a result of the large sample size and relatively low dimension of variables. However, we observed that many peaks clustered into one almost exact point on the PC plot. This is likely because they contain same types of motifs but different counts. According to the “TF collective” model of enhancer grammar, numbers of each TF-binding motif in an enhancer exhibit certain degrees of flexibility for its activity.^28^ A high-throughput screening of CRE activities of motifs showed that most motifs exhibit relatively low activity as a single functional unit of a CRE, and their activity linearly increase as the number increases.^29^ We thus assume that the combination of motifs plays a more important role than the number of each type of motifs, and then focused on the presence or absence of each motif and excluded their counts. The motif count matrices thus can be collapsed into binary-entry matrices that indicate if the motif exists (“1”) in a peak or not (“0”). This brings the 49,000 to 79,000 peaks in different treatment groups down to around 1064 “CRE types” defined by their motif components, which is less than the total possible combinations of the 11 motifs (2048 theoretical ways of combinations).

Generally, each STARR-seq-defined CRE type contains at least 3 different motifs, which is consistent with the fact that eukaryotic CREs are usually controlled by multiple factors. We defined combination-repressed CRE types in the DMSO group as types of more than a two-fold reduction of normalized reads from all peaks that belong to that type. We found 188 types out of over 1000 that are repressed (Figure 5c). Additionally, most of the repressed types contain E-boxes (93/188), confirming our hypothesis that the E-box is one of the major active motifs that is repressed by the drug combination. We also trained a logistic model based on single motifs and found that IRF, E2F, EBOX, and ETS are among the most important contributors to the repression by the drug combination (Supplementary Figure 8d). In all, the repressed motifs may represent a transcriptional mechanism that is yet to be revealed.

To understand more specific features of the repressed CRE types, we visualized motif distributions in the peaks that belong to them by finding motif clusters.^30^ The numbers and locations of the motifs show great diversity, and no obvious unifying feature were noticed, but they generally match the defined types. Interestingly, Most E-boxes accompany another motif in proximity (<50 bp, Figure 5c), implying that they might collaborate functionally, such as in DNA binding. To quantify the potential effect of motif combinations on the responsiveness to drug-combination treatment, for each CRE we constructed a vector that indicates if each pair of motifs cooccur in this CRE and performed logistic regression (Figure 5d). All the pairs that lost co-occurrence significance in Fisher’s test all appeared as positive contributors in logistic regression, confirming the consistency of our result (Figure 5d, red points). This logistic model can correctly classify 87% of the defined repressed type, but also shows a relatively high rate of false positives (50%). A more focused library will be needed to define a more refined model to quantify CRE types that are potentially repressed by the drug combination.

In summary, by using unbiased high-throughput CRE screening, we provided evidence that combining a CRY stabilizer with proteasome inhibitors suppresses the circadian gene transcription program by inhibiting CREs of specific types defined by their distinct combinations of TFBSs. Although more focused screenings and functional assays need to be done to validate and quantify the function of these CRE types, the results laid a foundation for the concept that CREs can be manipulated by small molecules and serve as a potential therapeutic modality in TNBC.

## Discussion

Genes that control the circadian rhythm have emerged as important regulators of cancer, and the growing availability of their small molecule modulators makes them attractive targets for cancer treatment. However, the complexity of their biological functions in cancer has only started to be revealed. As ubiquitous regulators of cell activity, circadian genes can become oncogenic^10^ or tumor-suppressing^31^ in different tumor types and tissue contexts. Previously we reported that in glioblastoma stem cells, BMAL1 has an altered binding landscape in the genome compared to noncancerous cells, resulting in new binding sites that rewire the metabolic gene networks to fuel tumor growth and also drive stemness.^10^ Others also reported the important roles of CLOCK in regulating the tumor microenvironment and immune responses in glioblastoma.^32^ However, a deeper biological model that connects their direct function, transcription, to physiological activities in cancer requires elaboration.

Breast cancer is among the types of cancer that are better understood in terms of their underlying biology and potential treatment options. To provide comparative insights, we selected breast cancer as our model for investigating the oncogenic function of BMAL1 and CLOCK, with a particular focus on the transcriptional activity, using small molecule modulators as experimental tools. We first showed that circadian genes have clinical relevance in the diagnosis and prognosis of breast cancer patients. Then we used RNAi to screen TNBC cell lines that are derived from different molecular subtypes. We found that only mMSL cells rely on BMAL1 and CLOCK to proliferate. This result reinforced our current knowledge that cancer cells with highly stem cell-like features may rely on BMAL1 and CLOCK.^9,10^ We thus applied small molecules to target their transcriptional activity in these TNBC cells.

Transcription factors are usually small and contain mostly intrinsically disordered regions, thus it is extremely challenging to find selective small molecule ligands to bind them and manipulate their functions. However, in cases where effective drugs are available, they exemplify cancer therapies with the best outcomes. For example, antagonizing estrogen receptor (ER), which is a ligand-binding transcription factor, can achieve a 90% five-year survival in ER-positive breast cancer.^14,33^ Inhibiting HIF2A, a TF that belongs to the same b-HLH-PAS family as BMAL1 and CLOCK, with the small molecule drug Belzutifan can effectively suppress multiple types of tumors associated with the von Hippel-Lindau disease.^34^ Here we leveraged the negative arms of the circadian network to indirectly achieve repression of BMAL1 and CLOCK activity. Probably due to the core circadian network is rewired in these highly malignant TNBC cells, only the CRY stabilizer SHP1705 achieved faithful repression of core BMAL1 and CLOCK targets. To expand the repressive effect of SHP1705, we added inhibitors of the proteasome, which was previously reported to be indispensable in maintaining BMAL1 and CLOCK-mediated transcription. This drug combination indeed displayed synergy against cell proliferation and effectively repressed circadian cycling genes.

This synergy provides us both a leverage to study the transcriptional programs for which BMAL1 and CLOCK are responsible in fueling tumor growth, and a potential therapeutic strategy to suppress that program. In ER-driven breast cancer, ER serves as the master TF that can strongly induce downstream transcriptional programs upon binding to its activating ligand.^35,36^ However. in mMSL TNBC, BMAL1 and CLOCK do not appear to function as a “binary switch” as ER, because a CRY stabilizer faithfully repressed well-established B/C target genes but has little impact on the global transcriptome and failed to recapitulate the strong phenotype observed following genetic knockdown of B/C.

To understand the disparity between these driver TFs of breast cancer, we tried to examine their function from the DNA side, that is, the CREs through which the TFs function. ER’s potent activity is mediated by the strong enhancer activities induced by it to drive transcription.^37^ These enhancers that are activated by singular kinds of factors are common among developmental and lineage-determining TFs, but the similar concepts are less applicable for constitutive housekeeping TFs like BMAL1 and CLOCK.^38,39^ In an analysis of super enhancers (SEs) in TNBC, a group of TF binding motifs including E-box was identified to constitute the SEs.^23^ Although currently we have no evidence to support that this is a SE-mediated mechanism, this finding inspired us to hypothesize that strong CREs in TNBC can be activated and maintained by multiple TFs in the absence of a singular master driver.

Following this logic, we examined CRE activities after drug treatment and defined features of CREs that are potentially repressed by the combination of CRY2 stabilizer and proteasome inhibitor. These CREs comprise different combinations of the binding motifs of several constitutive TFs, such as SP1, E2Fs, NFYs, and YY1. These factors may collaboratively activate CREs that drive tumor-fueling transcription programs. The mechanism by which the effects of small molecules are translated from these *trans*-acting (TFs) to *cis*-acting factors remains to be studied. We hypothesize that it is achieved by interfering with the dynamics of the CREs since both drugs disrupt the turnover of the TFs bound to the CREs. Nonetheless, our results revealed a potential mechanism by which we can target the activity of selective CREs based on their regulatory properties. This model provided a new perspective of modulating gene transcription programs with higher precision in cancer, opening a door for defining cancer drivers and biomarkers based on *cis*-regulatory programs, and tailoring corresponding therapies.

Meanwhile, this model is still primitive because of several limiting factors. First, our current library lacks sufficient focus to provide a more detailed and determinative description of the features of CREs that are repressed by the drugs. Second, dependencies of the active CREs on core promoter features cannot be assayed by the current STARR-seq setup because the vector uses a TATA-box-dependent promoter, whereas E-box-containing CREs also often work with CG-rich promoters that lack a TATA-box.^40^ Third, there is a deficiency in both our knowledge of the intricate biology, especially the dynamics, of CREs and the corresponding computational methodologies. In the future, more comprehensive quantifications of CRE features, such as motif composition and position dependency, needs to be done with more focused library screening and more precise computational methods to better define the target CREs of the drugs. Mechanistic studies are also needed to strengthen the biological rationale for targeting *cis*-element activities through manipulating their partner *trans*-acting factors. Finally, to translate this model into pre-clinical and clinical studies, optimizing proteasome inhibitors for in vivo stability might be required. This optimization should aim to maintain the global proteostasis of cells while effectively targeting transcription. In summary, this study identified the master circadian clock genes BMAL1 and CLOCK as potential targets in the mMSL type of TNBC. Furthermore, we proposed a new biological model for targeting CREs with small molecules. These results offer new opportunities for understanding driving mechanisms, discovering new biomarkers, and designing therapeutics for TNBC.

## Methods

### TCGA and METABRIC data analysis

TCGA-BRCA gene expression and clinical data were downloaded from UCSC Xena.^41^ METABRIC cohort data were downloaded from cBioportal.^42^ DE genes analysis between tumor and normal tissues were defined using Wilcoxon rank test with FDR < 0.05.43 Logistic regression was done using the R package `glm`. Multi-variate Cox regression was done using the R package `coxph`.

### Cell Culture and shRNA screening

All cell lines are obtained from ATCC. MDA-MB cell lines are cultured in RPMI-1640 (Gibco cat. 72400-074) containing 10% Fetal Bovine Serum (FBS, RnD Systems cat. S11150), all other cell lines are cultured per ATCC’s recommended media. All cells are cultured in a 37 °C incubator with 5% CO2. shRNA-vectors are produced as described in our previous work.^10^ Briefly, lentivirus containing shRNA was transfected into HEK 293T cells using Lipofectamine 3000 (Invitrogen cat. L3000015) reagents per manufacturer’s manual, and the supernatant containing lentivirus was harvested 24 hours after transfection.

For proliferation screening after shRNA KD, cells were plated in 6-well plates to reach a 30-50% confluency by the time of transduction. Lentiviruses were diluted in culture media containing 5µg/mL Polybrene (Sigma cat. TR-1003-G) to the same MOI and added to cells. 48 hours after transduction, KD efficiency and puromycin selection were performed. To validate KD, RNA was extracted and reverse-transcribed into cDNA to run qPCR as described in the following section. To select for KD cells, media was exchanged into fresh culture media containing 2 µg/mL puromycin. 72 hours after puromycin addition, cells were stained with crystal violet. The screening was repeated three times.

For quantifying the proliferation of mMSL cell lines after KD, lentiviral transduction was performed. After forty-eight hours, cells were treated with 2µg/mL puromycin for another 48 hours, then plated into black clear-bottom 96-well plates (Falcon cat. 353219) 48 hours after transduction in a density of 10,000 cells per well. At indicated time points (day 0, 2, and 4), 50µL CellTiter-Glo (Promega cat. G7572) was added to each well, incubated for 15 minutes at room temperature, and read in a TECAN luminescence reader. Proliferation was normalized to day 0.

### EdU assay

EdU assay was performed with the Invitrogen Click-iT Plus EdU Pacific Blue kit (Invitrogen cat # C10636) following manufacturer’s protocol. Briefly, after 48 hours of shRNA transduction, EdU (final concentration 10 µM) was added in cultured cells and incubated at 37°C for one hour. The cells were then harvested with TrypLE, washed with 3mL 1% BSA in PBS, and fixed with 100 µL component D at room temperature for 15 mins. Wash the cells with 3mL 1% BSA in PBS and resuspend in 3 mL 100 µL of 1X permeabilization and wash reagent, incubate for 15 minutes. The click reaction cocktail was freshly prepared before the experiment, and 500 µL was added to each sample of fixed cells and mixed well. The reaction was incubated for 30 mins at room temperature, then washed once with 1X permeabilization and wash reagent before analyzing. The samples were analyzed on a Invitrogen Attune NxT flow cytometer.

### Quantitative RT-PCR

Total RNA was extracted with the NEB Monarch RNA miniprep kit (NEB cat. T2010S) following manufacturer’s manual with on-column DNA digestion. 500 ng total RNA from each sample was used to perform reverse transcription using SuperScript IV VILO Master Mix (Invitrogen cat. 11756050) following manufacturer’s manual. Real-time PCR was done using PowerUp SYBR Green Master Mix (Applied Biosystems cat. A25742) in a thermal cycler (Bio-Rad). Cq value was defined using regression, normalized to PPIA^44^, and transformed to 2^ΔΔCq^.

### Small molecule screening and synergy analysis

Cells were seeded in clear-bottom black 96-well plates (10,000 cells per well). Twenty-four hours after seeding, fresh media was added in each well and small molecules of indicated concentration were added via two-fold serial dilution. CellTiter-Glo (Promega cat. G7572) was added to each well 72 hours after the addition of small molecules. The plate was incubated at room temperature for 15 minutes and luminescence was determined in a TECAN plate reader. All small molecule experiments were repeated at least three times. Synergistic effects were quantified using the *SyngergyFinder* software^45^.

### RNA sequencing

Total RNA was extracted using NEB Monarch Total RNA miniprep kit (NEB cat. T2010S) following manufacturer’s manual with on-column DNase I digestion. For shRNA KD experiments, total RNA was extracted 48 hours after transduction. For small molecules-treated cells, total RNA was extracted 8 or 24 hours after adding small molecules. mRNA was selected through poly-A tail selection, library preparation and sequencing were done with *Azenta*.

### RNA-seq data analysis

Pair-end reads in FASTQ format were processed with `trimmomatic`^46^ or `trimgalore`^47^ to trim off adaptor sequences, only paired reads were kept, and sequencing quality was checked with `fastqc`^48^ and `multiqc`^49^. QC-passed data were aligned to human genome *hg38* with `hisat2`^50^ and gene counts were summarized using `featureCounts`^51^. Differential gene expression was done with `DEseq2`^52^. GSEA was performed using the GSEA software^53^ or with `fgsea`^54^, DE genes were ranked by log2FoldChange and pre-rank GSEA analysis were performed with 1000 permutations. GO and TF enrichment was performed using `gProfiler2`^55^ and `enrichr`^56^. Volcano plots of DE genes are plotted using the R package `EnhancedVolcano57`.

### MDA-MB-231 H3K27ac ChIP-seq analysis

Raw sequencing reads were downloaded from GEO (GSE85158)^58^. Following trimming and quality control using `trimmomatic` and `fastqc` respectively, reads were aligned to human genome hg38 using bowtie2^59^, and broad peaks were called using MACS2^60^. Motifs in the peaks were counted using the `countPWM` function from the `Biostrings`^61^ package in R with matching threshold set to 87%. Co-occurrence test was performed using Fisher’s exact test in R, and the p values were corrected using the Bonferroni method.

### Cancer Cell Line Encyclopedia (CCLE) data analysis

Gene expression data of CCLE lines were downloaded from the Broad Institute DepMap portal (https://depmap.org/portal/download/all/).^24^ Correlation analyses were implemented and plotted using the corrplot package in R.

### Luciferase reporter assay

Promoters of genes of interest were cloned from the genomic DNA of MDA-MB-231 cells and cloned into an EcoRI-linearized luciferase reporter vector using In-Fusion cloning (Takara Bio cat. 638948). Lentivirus was packaged using Lipofectamine 3000 following manufacturer’s instructions. The reporter cell lines were established by lentiviral transduction followed by 10 µg/mL blasticidin (Gibco cat. A1113902) selection for 96 hours. For single time point luciferase assay, cells were cultured for 8 hours after small molecules were added and BrightGlo (Promega cat. G2650) luciferase assay substrate was added to each well and luminescence was read in a TECAN plate reader. For prolonged luciferase activity recording, small molecules were added in circadian luciferase assay media, which was added in cells, and luminescence was recorded every hour for the indicated time length in a TECAN plate reader. Triplicates were performed for each condition and mean values were plotted.

### STARR-seq library preparation and screening

STARR-seq screening was performed as described by Muerdter and Boryn *et al.* with minor adjustments^62^. Briefly, the genomic library (700 bp on average) was prepared by sonicating genomic DNA extracted from MDA-MB-231 cells, adaptor-ligated using NEBNext Ultra II DNA library preparation kit for Illumina, and expanded by PCR using Roche KAPA HiFi HotStart ReadyMix (Roche cat. KK2602). hSTARR-ORI vector (Addgene #99296) was linearized by dual restriction enzyme digestion with AgeI-HF (NEB cat. R3552S) and SalI-HF (NEB cat. R3138T). The linearization reaction was run on 1% agarose gel, the 3000bp band was cut out, and DNA was extracted using Nucleospin Gel and PCR Clean-up kit (Macherey-Nagel Cat. 740609) and cleaned with QIAGEN MinElute PCR purification kit (QIAGEN cat. 28006). The library was inserted into the vector using In-Fusion cloning. The resulting vectors were transformed into MegaX DH10B electrocompetent cells (Invitrogen cat. C640003) with GenePulser (Bio-rad). Transformed cells were recovered in 1mL recovery media at 37°C for one hour and cultured overnight in 12L LB media. Library-containing plasmids were extracted using QIAGEN Plasmid Plus Giga Kit (QIAGEN cat. 12991) following manufacturer’s instructions.

The STARR-screening library was transformed into MDA-MB-231 cells containing small molecules using Lipofectamine 3000. Total RNA was extracted six hours after transformation using QIAGEN RNeasy Maxi Kit (QIAGEN cat. 75162) following manufacturer’s manual. Poly-A+ RNA was selected using Dynabeads Oligo(dT)25 (Invitrogen cat. 61005) and reverse transcription was performed using SuperScript III (Invitrogen cat. 18080093) with gene-specific primers (GSP). Library preparation for sequencing was performed using a two-step amplification (junction PCR followed by sequencing-ready PCR) and cleaned using SPRIselect beads (Beckman cat. B23318). Sequencing was done with *Azenta*.

### STARR-seq data analysis

After adaptor trimming and quality control using `trimmomatic` and `fastqc` respectively, reads were aligned to the human genome *hg38* using `bowtie2`.^59^ Peak calling was done using `MACS263`. Motif counting and co-occurrence analyses were done as described in the analysis for H3K27ac ChIP-seq data. Motif clusters were found and visualized using `MCAST64`.with the following parameters: `mcast –hardmask --output-ethresh 10.0 --max-gap 200 --max-total-width 1000 --motif-pthresh 5.0E-4`. Position Fraction Matrices (PFMs) for clustered TFBS were downloaded from JASPAR 2022 (https://jaspar2022.genereg.net/downloads/)^27^. The PFM for SIX5 was obtained from geneXplain database (https://genexplain.com).

### Statistical Analysis

Log-rank test was performed for all K-M curves using `ggsurvplot`. P-values for qPCR and CellTiter-Glo were calculated by one-way ANOVA test. P-values for luciferase reporter assay for promoter activities are calculated using unpaired t-test. All other statistical methods are specified in corresponding method sections.

### Data and code availability

All newly generated raw sequencing data are available on GEO (GSE266013). All codes and scripts used to analyze the data are available on GitHub (https://github.com/yzpan1/targeting-clock-in-tnbc/tree/main).

### Supplementary Methods

Detailed descriptions of methods, oligo sequences, and data analyses are available in supplementary methods.

## Authors’ Contributions

Y.P. and S.A.K. conceptualized and designed the project. Y.P. designed and performed experiments, acquired data, and led data analysis. Y.P., L.Z., P.C., T.T.K., F.B., S.S., P.J., H.J.L, S.M.M., E.R.T., and S.A.K performed data analysis and discussion. F.B., P.J., H.J.L., and E.R.T. provided consultation on clinical data analysis. Y.P., T.P.C., and R.R. performed quantitative motif analysis. J.J.L supervised all statistical methods. Y.P., E.R.T., and S.A.K. wrote the manuscript with input from all authors. E.R.T. and S.A.K. supervised the project.

## Acknowledgments

We appreciate the Translational Team Accelerator Program funding to Steve A. Kay, Heinz-Josef Lenz, and Evanthia Roussos Torres. We also thank the Center for Advanced Research Computing of USC. This work was also supported by the National Institutes of Health (grant R35GM130376 to R.R.).

## References

1. Takahashi, J. S. Transcriptional architecture of the mammalian circadian clock. Nat Rev Genet 18, 164–179 (2017).

2. Allada, R. & Bass, J. Circadian Mechanisms in Medicine. New England Journal of Medicine 384, (2021).

3. Patke, A., Young, M. W. & Axelrod, S. Molecular mechanisms and physiological importance of circadian rhythms. Nature Reviews Molecular Cell Biology vol. 21 Preprint at 10.1038/s41580-019-0179-2 (2020).

4. Mure, L. S. et al. Diurnal transcriptome atlas of a primate across major neural and peripheral tissues. Science *(1979)* 359, (2018).

5. Sulli, G., Lam, M. T. Y. & Panda, S. Interplay between Circadian Clock and Cancer: New Frontiers for Cancer Treatment. Trends Cancer 5, 475–494 (2019).

6. Battaglin, F. et al. Clocking cancer: the circadian clock as a target in cancer therapy. Oncogene 40, 3187–3200 (2021).

7. Pariollaud, M. & Lamia, K. A. Cancer in the Fourth Dimension: What Is the Impact of Circadian Disruption? Cancer Discov 10, 1455–1464 (2020).

8. Masri, S. & Sassone-Corsi, P. The emerging link between cancer, metabolism, and circadian rhythms. Nat Med 24, 1795–1803 (2018).

9. Puram, R. V. et al. Core Circadian Clock Genes Regulate Leukemia Stem Cells in AML. Cell 165, 303–316 (2016).

10. Dong, Z. et al. Targeting glioblastoma stem cells through disruption of the circadian clock. Cancer Discov 9, 1556–1573 (2019).

11. Curtis, C. et al. The genomic and transcriptomic architecture of 2,000 breast tumours reveals novel subgroups. Nature 486, (2012).

12. Li, S. Y. et al. Tumor circadian clock strength influences metastatic potential and predicts patient prognosis in luminal A breast cancer. Proc Natl Acad Sci U S A 121, (2024).

13. Anafi, R. C., Francey, L. J., Hogenesch, J. B. & Kim, J. CYCLOPS reveals human transcriptional rhythms in health and disease. Proc Natl Acad Sci U S A 114, 5312–5317 (2017).

14. Waks, A. G. & Winer, E. P. Breast Cancer Treatment: A Review. JAMA - Journal of the American Medical Association 321, 288–300 (2019).

15. Lehmann, B. D. et al. Identification of human triple-negative breast cancer subtypes and preclinical models for selection of targeted therapies. Journal of Clinical Investigation 121, (2011).

16. He, Y. et al. Structure-Activity Relationship and Biological Investigation of a REV-ERBα-Selective Agonist SR-29065 (34) for Autoimmune Disorders. J Med Chem 66, 14815–14823 (2023).

17. Oshima, T. et al. Cell-based screen identifies a new potent and highly selective CK2 inhibitor for modulation of circadian rhythms and cancer cell growth. Sci Adv 5, 1–16 (2019).

18. Humphries, P. S. et al. Carbazole-containing amides and ureas: Discovery of cryptochrome modulators as antihyperglycemic agents. Bioorg Med Chem Lett 28, (2018).

19. Hirota, T. et al. Identification of small molecule activators of cryptochrome. Science *(1979)* 337, 1094–1097 (2012).

20. Stratmann, M., Suter, D. M., Molina, N., Naef, F. & Schibler, U. Circadian Dbp Transcription Relies on Highly Dynamic BMAL1-CLOCK Interaction with E Boxes and Requires the Proteasome. Mol Cell 48, 277–287 (2012).

21. Goldberg, A. L. Development of proteasome inhibitors as research tools and cancer drugs. Journal of Cell Biology 199, (2012).

22. Bozek, K. et al. Regulation of clock-controlled genes in mammals. PLoS One 4, e4882 (2009).

23. Huang, H. et al. Defining super-enhancer landscape in triple-negative breast cancer by multiomic profiling. Nat Commun 12, (2021).

24. Barretina, J. et al. The Cancer Cell Line Encyclopedia enables predictive modelling of anticancer drug sensitivity. Nature 483, (2012).

25. Pan, Y., van der Watt, P. J. & Kay, S. A. E-box binding transcription factors in cancer. Frontiers in Oncology vol. 13 Preprint at 10.3389/fonc.2023.1223208 (2023).

26. Arnold, C. D. et al. Genome-wide quantitative enhancer activity maps identified by STARR-seq. Science *(1979)* 339, (2013).

27. Castro-Mondragon, J. A. et al. JASPAR 2022: The 9th release of the open-access database of transcription factor binding profiles. Nucleic Acids Res 50, (2022).

28. Smith, G. D., Ching, W. H., Cornejo-Páramo, P. & Wong, E. S. Decoding enhancer complexity with machine learning and high-throughput discovery. Genome Biology vol. 24 Preprint at 10.1186/s13059-023-02955-4 (2023).

29. Sahu, B. et al. Sequence determinants of human gene regulatory elements. Nat Genet 54, (2022).

30. Andersson, R. & Sandelin, A. Determinants of enhancer and promoter activities of regulatory elements. Nat Rev Genet 21, 71–87 (2020).

31. Sha, A. A. et al. The circadian cryptochrome, CRY1, is a pro-tumorigenic factor that rhythmically modulates DNA repair. Nat Commun 12, 401 (2021).

32. Chen, P. et al. Circadian regulator CLOCK recruits immune-suppressive microglia into the GBM tumor microenvironment. Cancer Discov 10, 371–381 (2020).

33. Harbeck, N. & Gnant, M. Breast cancer. The Lancet 389, 1134–1150 (2017).

34. Fallah, J. et al. FDA Approval Summary: Belzutifan for von Hippel-Lindau Disease–Associated Tumors. Clinical Cancer Research 28, (2022).

35. Eeckhoute, J., Carroll, J. S., Geistlinger, T. R., Torres-Arzayus, M. I. & Brown, M. A cell-type-specific transcriptional network required for estrogen regulation of cyclin D1 and cell cycle progression in breast cancer. Genes Dev 20, (2006).

36. Deroo, B. J. & Korach, K. S. Estrogen receptors and human disease. Journal of Clinical Investigation vol. 116 Preprint at 10.1172/JCI27987 (2006).

37. Liu, Z. et al. Enhancer activation requires trans-recruitment of a mega transcription factor complex. Cell 159, (2014).

38. Dejosez, M. et al. Regulatory architecture of housekeeping genes is driven by promoter assemblies. Cell Rep 42, (2023).

39. Zabidi, M. A. et al. Enhancer-core-promoter specificity separates developmental and housekeeping gene regulation. Nature 518, (2015).

40. Lin, J. M. et al. Transcription factor binding and modified histones in human bidirectional promoters. Genome Res 17, 818–827 (2007).

41. Goldman, M. J. et al. Visualizing and interpreting cancer genomics data via the Xena platform. Nature Biotechnology vol. 38 Preprint at 10.1038/s41587-020-0546-8 (2020).

42. Cerami, E. et al. The cBio Cancer Genomics Portal: An open platform for exploring multidimensional cancer genomics data. Cancer Discov 2, (2012).

43. Li, Y., Ge, X., Peng, F., Li, W. & Li, J. J. Exaggerated false positives by popular differential expression methods when analyzing human population samples. Genome Biol 23, (2022).

44. Nakao, R., Okauchi, H., Hashimoto, C., Wada, N. & Oishi, K. Determination of reference genes that are independent of feeding rhythms for circadian studies of mouse metabolic tissues. Mol Genet Metab 121, (2017).

45. Ianevski, A., Giri, A. K. & Aittokallio, T. SynergyFinder 2.0: Visual analytics of multi-drug combination synergies. Nucleic Acids Res 48, (2021).

46. Bolger, A. M., Lohse, M. & Usadel, B. Trimmomatic: A flexible trimmer for Illumina sequence data. Bioinformatics 30, (2014).

47. Krueger, F. Trim Galore!: A wrapper tool around Cutadapt and FastQC to consistently apply quality and adapter trimming to FastQ files. Babraham Institute (2015).

48. Andrews, S. FastQC - A quality control tool for high throughput sequence data. http://www.bioinformatics.babraham.ac.uk/projects/fastqc/. Babraham Bioinformatics (2010).

49. Ewels, P., Magnusson, M., Lundin, S. & Käller, M. MultiQC: Summarize analysis results for multiple tools and samples in a single report. Bioinformatics 32, (2016).

50. Kim, D., Paggi, J. M., Park, C., Bennett, C. & Salzberg, S. L. Graph-based genome alignment and genotyping with HISAT2 and HISAT-genotype. Nat Biotechnol 37, (2019).

51. Liao, Y., Smyth, G. K. & Shi, W. FeatureCounts: An efficient general purpose program for assigning sequence reads to genomic features. Bioinformatics 30, (2014).

52. Love, M. I., Huber, W. & Anders, S. Moderated estimation of fold change and dispersion for RNA-seq data with DESeq2. Genome Biol 15, (2014).

53. Subramanian, A. et al. Gene set enrichment analysis: A knowledge-based approach for interpreting genome-wide expression profiles. Proc Natl Acad Sci U S A 102, (2005).

54. Korotkevich, G., Sukhov, V., Budin, N., Atryomov, M. N. & Sergushichev, A. Fast gene set enrichment analysis. bioRxiv. bioRxiv (2021).

55. Peterson, H., Kolberg, L., Raudvere, U., Kuzmin, I. & Vilo, J. gprofiler2 -- an R package for gene list functional enrichment analysis and namespace conversion toolset g: Profiler. F1000Res **9**, (2020).

56. Kuleshov, M. V. et al. Enrichr: a comprehensive gene set enrichment analysis web server 2016 update. Nucleic Acids Res 44, (2016).

57. Blighe K, Rana S & Lewis M. Publication-ready volcano plots with enhanced colouring and labeling. Bioconductor (2022).

58. Franco, H. L. et al. Enhancer transcription reveals subtype-specific gene expression programs controlling breast cancer pathogenesis. Genome Res 28, (2018).

59. Langmead, B. & Salzberg, S. L. Fast gapped-read alignment with Bowtie 2. Nat Methods 9, (2012).

60. Zhang, Y. et al. Model-based analysis of ChIP-Seq (MACS). Genome Biol 9, (2008).

61. Pagés, H., Aboyoun, P., Gentleman, R. & DebRoy, S. Biostrings: Efficient manipulation of biological strings. R package version 2.58.*0*. Preprint at (2020).

62. Muerdter, F. et al. Resolving systematic errors in widely used enhancer activity assays in human cells. Nat Methods 15, (2018).

63. 63. Gaspar, J. M. Improved peak-calling with MACS2. bioRxiv (2018).

64. Bailey, T. L. & Noble, W. S. Searching for statistically significant regulatory modules. in Bioinformatics vol. 19 (2003).

