## Supplementary Figure, Methods, and Materials for "Targeting circadian transcriptional programs through a cis-regulatory mechanism in triple negative breast cancer"

**Supplementary Figure 1. a.** Negative correlation between ARNTL and negative arm genes are lost in tumor tissues in both TCGA and METABRIC cohort. **b.** Differential expression analysis of core circadian clock genes in TCGA cohort. Log2-fold change was plotted, and significance level was based on Wilcoxon test. **c-d.** Significant results of using the median expression level of single circadian genes to stratify patient cohorts and test difference on overall survival in either all patients (c) or in different subtypes (d). **e.** Logistic model that can separate tumor and normal tissue at an over 90% accuracy has no implications on survival for TCGA cohort (left). The model produce a statistically significant result on median survival time for the METABRIC cohort (right), but survival time cannot be separated towards the end of disease. **f.** Validation of multivariate Cox-regression model on the TCGA cohort data.

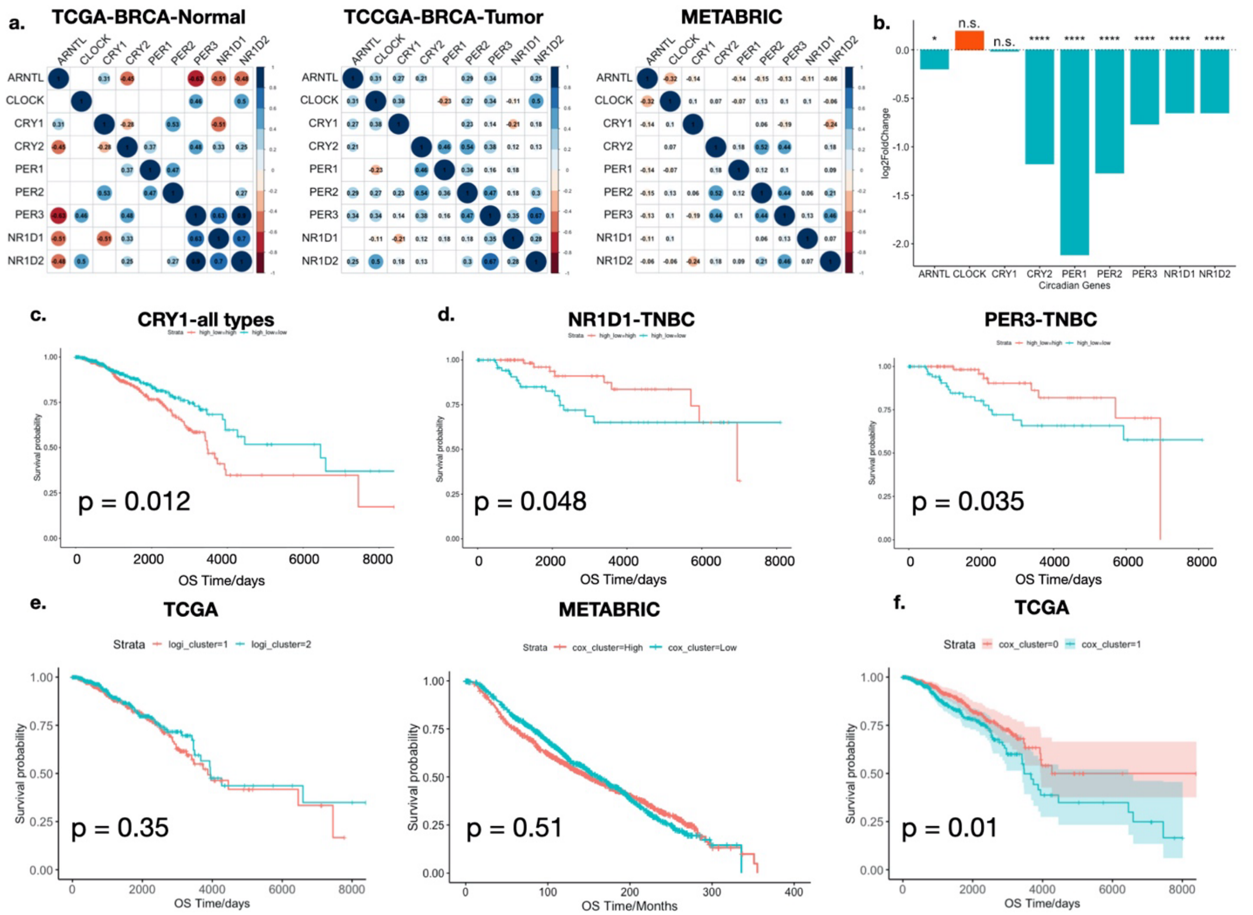

**Supplementary Figure 2. a.** The TNBC cell line panel screened for BMAL1 and CLOCK knockdown. MSL: Mesenchymal Stem-like; LAR: Luminal androgen receptor; M: Mesenchymal; BL: Basal-like. **b.** Initial gene knockdown (KD) screening of TNBC cell lines. Cell proliferation after KD was visualized by crystal violet staining (left). KD efficiency was validated using quantitative RT-PCR (right).

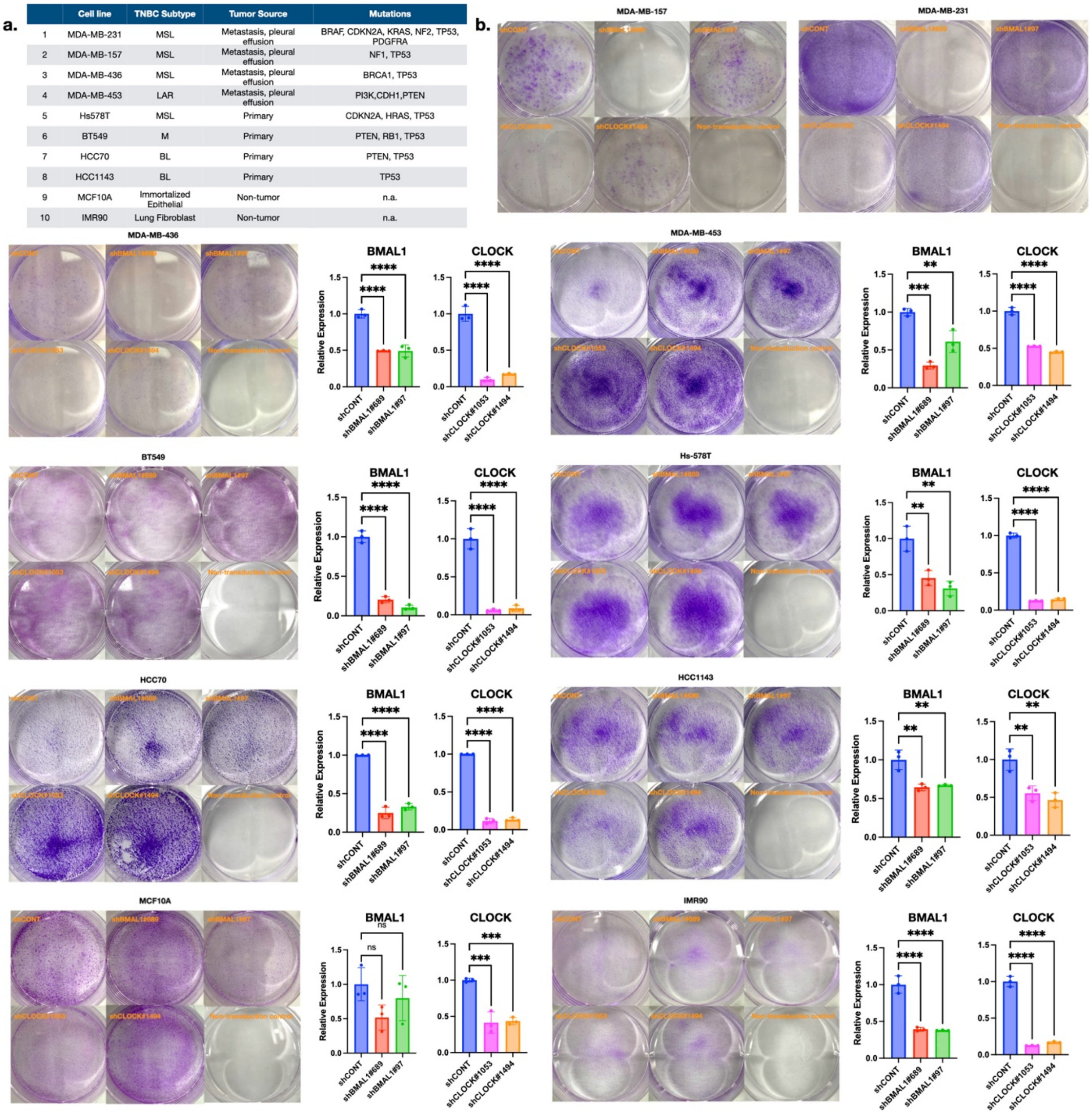

| | Cell line | TNBC Subtype | Tumor Source | Mutations |
|---|---|---|---|---|
| 1 | MDA-MB-231 | MSL | Metastasis, pleural effusion | BRAF, CDKN2A, KRAS, NF2, TP53, PDGFRA |
| 2 | MDA-MB-157 | MSL | Metastasis, pleural effusion | NF1, TP53 |
| 3 | MDA-MB-436 | MSL | Metastasis, pleural effusion | BRCA1, TP53 |
| 4 | MDA-MB-453 | LAR | Metastasis, pleural effusion | PI3K,CDH1,PTEN |
| 5 | Hs578T | MSL | Primary | CDKN2A, HRAS, TP53 |
| 6 | BT549 | M | Primary | PTEN, RB1, TP53 |
| 7 | HCC70 | BL | Primary | PTEN, TP53 |
| 8 | HCC1143 | BL | Primary | TP53 |
| 9 | MCF10A | Immortalized Epithelial | Non-tumor | n.a. |
| 10 | IMR90 | Lung Fibroblast | Non-tumor | n.a. |

**Supplementary Figure 3. a.** Quantification of cell proliferation after BMAL1 and CLOCK knockdown using CellTiter. p-value comparing the relative proliferation on day 4 was calculated using one-way ANOVA. n=4. *: p<0.05; **: p<0.01; ***: p<0.001; ****: p<0.0001. **b.** Volcano plots summarizing DE genes after BMAL1 and CLOCK knockdown in MDA-MB-231 cells. **c.** EdU flow cytometry assay showing reduced proportion of EdU-positive cells indicating reduced cells going through cell cycle. p-value was calculated using one-way ANOVA. n=3. *: p<0.05; **: p<0.01; ***: p<0.001; ****: p<0.0001.

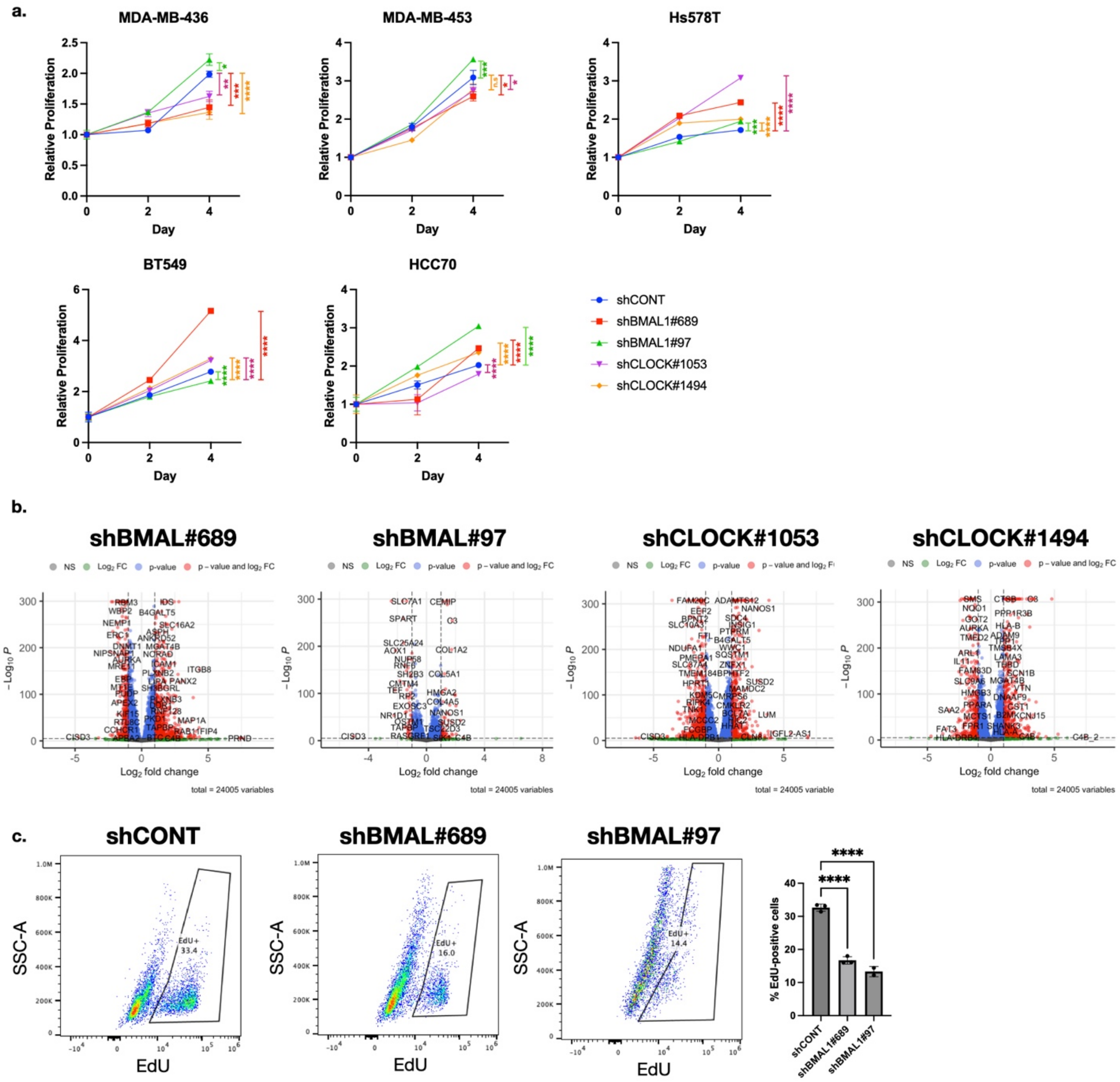

**Supplementary Figure 4. a.** Diagram of the small molecules that target the core circadian gene network to repress the activity of BMAL1 and CLOCK. CRY stabilizers prevent CRY from being ubiquitinated and degraded by the proteasome, thus prolong the repression of BMAL1 and CLOCK dimer activity. CK2 inhibitors achieve this through preventing the phosphorylation of PER proteins and eventually their degradation. REV-ERB agonists elevates REV-ERB proteins' repressive function on the transcription of BMAL1 gene. **b.** Volcano plot of DE genes after SHP1705 treatment for 24 hours. **c.** GSEA results showed that as a single agent, SHP1705 has negligible effect on the hallmark pathways, explaining its lack of effect on cell proliferation. **d.** Synergy curve and average Bliss synergy score of SHP1705 and carfilzomib in three mMSL TNBC cells.

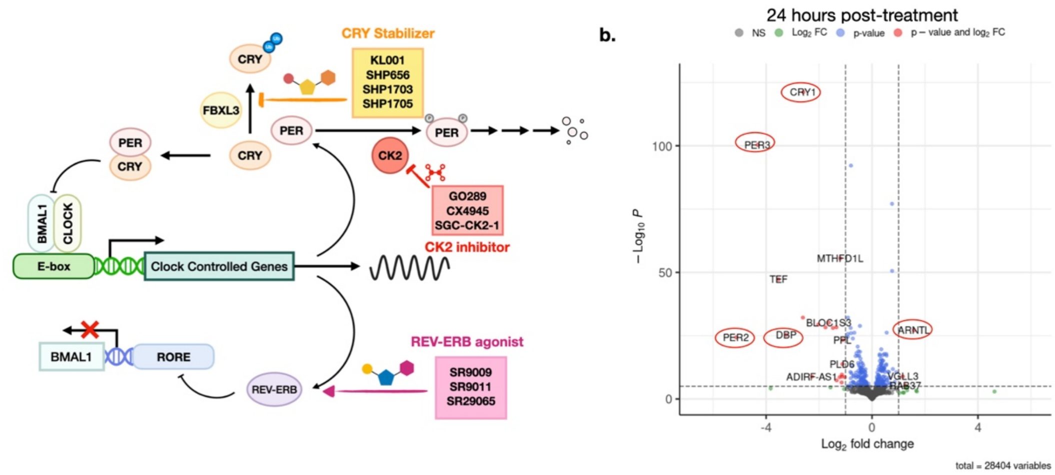


c.

| | GS follow link to MSigDB | GS DETAILS | SIZE | ES | NES | NOM p-val | FDR q-val | FWER p-val |
|---|---|---|---|---|---|---|---|---|
| 1 | HALLMARK_KRAS_SIGNALING_UP | Details ... | 190 | -0.47 | -1.60 | 0.000 | 0.241 | 0.116 |
| 2 | HALLMARK_EPITHELIAL_MESENCHYMAL_TRANSITION | Details ... | 194 | -0.48 | -1.55 | 0.000 | 0.244 | 0.264 |
| 3 | HALLMARK_INTERFERON_GAMMA_RESPONSE | Details ... | 192 | -0.34 | -1.55 | 0.000 | 0.182 | 0.264 |
| 4 | HALLMARK_INFLAMMATORY_RESPONSE | Details ... | 188 | -0.35 | -1.53 | 0.000 | 0.182 | 0.316 |
| 5 | HALLMARK_TNFA_SIGNALING_VIA_NFKB | Details ... | 198 | -0.29 | -1.51 | 0.000 | 0.220 | 0.363 |
| 6 | HALLMARK_HYPOXIA | Details ... | 197 | -0.26 | -1.51 | 0.000 | 0.209 | 0.404 |
| 7 | HALLMARK_IL6_JAK_STAT3_SIGNALING | Details ... | 79 | -0.41 | -1.46 | 0.000 | 0.211 | 0.518 |
| 8 | HALLMARK_APOPTOSIS | Details ... | 158 | -0.24 | -1.44 | 0.000 | 0.220 | 0.609 |
| 9 | HALLMARK_COAGULATION | Details ... | 129 | -0.35 | -1.41 | 0.000 | 0.286 | 0.661 |
| 10 | HALLMARK_APICAL_JUNCTION | Details ... | 189 | -0.32 | -1.40 | 0.000 | 0.268 | 0.661 |
| 11 | HALLMARK_INTERFERON_ALPHA_RESPONSE | Details ... | 97 | -0.37 | -1.40 | 0.075 | 0.249 | 0.661 |

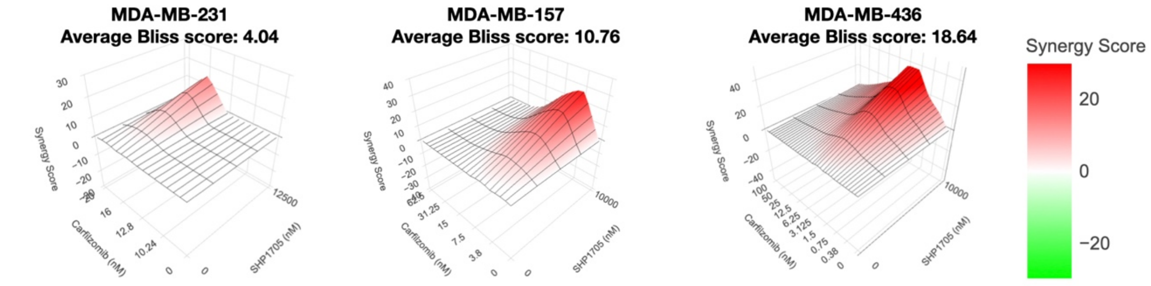

**Supplementary Figure 5.** a. Raw counts RNA-seq data showing that expression level change of selective core BMAL1 and CLOCK target genes is consistent with qPCR results after SHP1705 treatment. **b-c.** Volcano plot showing gene expression change after MG132 (b) or MG132 and SHP1705 combination (c) treatment. **d.** Enrichment analysis of transcription factors after SHP1705 treatment. **e.** Volcano plot showing significantly changed genes after carfilzomib treatment for 8 hours. **f.** Venn diagram showing that in the SHP1705 and carfilzomib combination, like in SHP1705 and MG132 treatment, there is a set of genes that are specific to the combination group that might account for the combination effect. **g.** Refined list of motifs can represent the binding sites of the enriched TFs. **h.** Positive correlation between β-subunit of EBTFs and EBTF-Co are observed in the tumor samples of the TCGA-BRCA database.

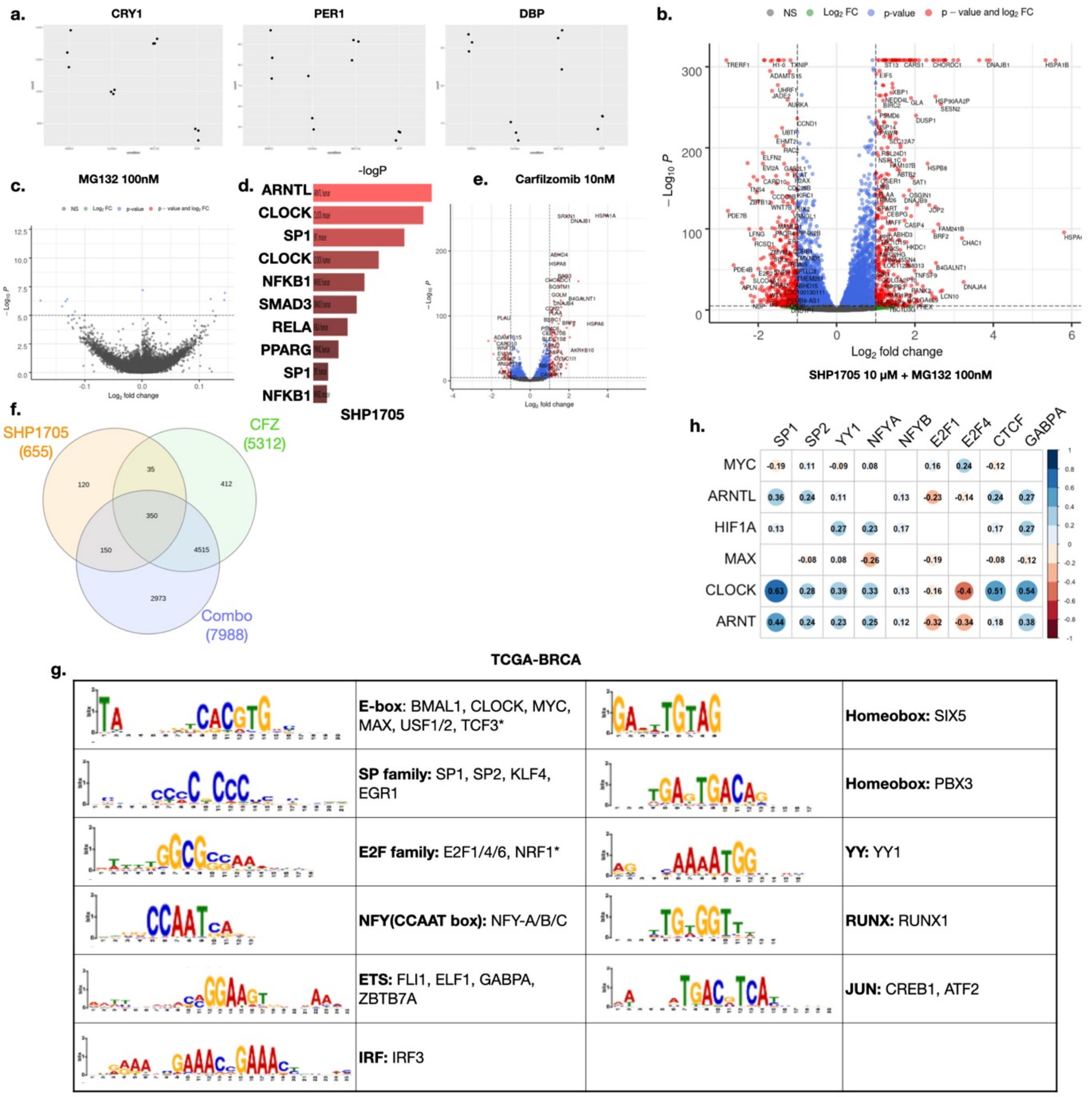

**Supplementary Figure 6. a.** Sequence and TFBS features of the promoter sequences that are tested by luciferase activity assay. Different combinations of EBTF-Co motifs are observed, suggesting their activity are likely to be repressed by the combination.

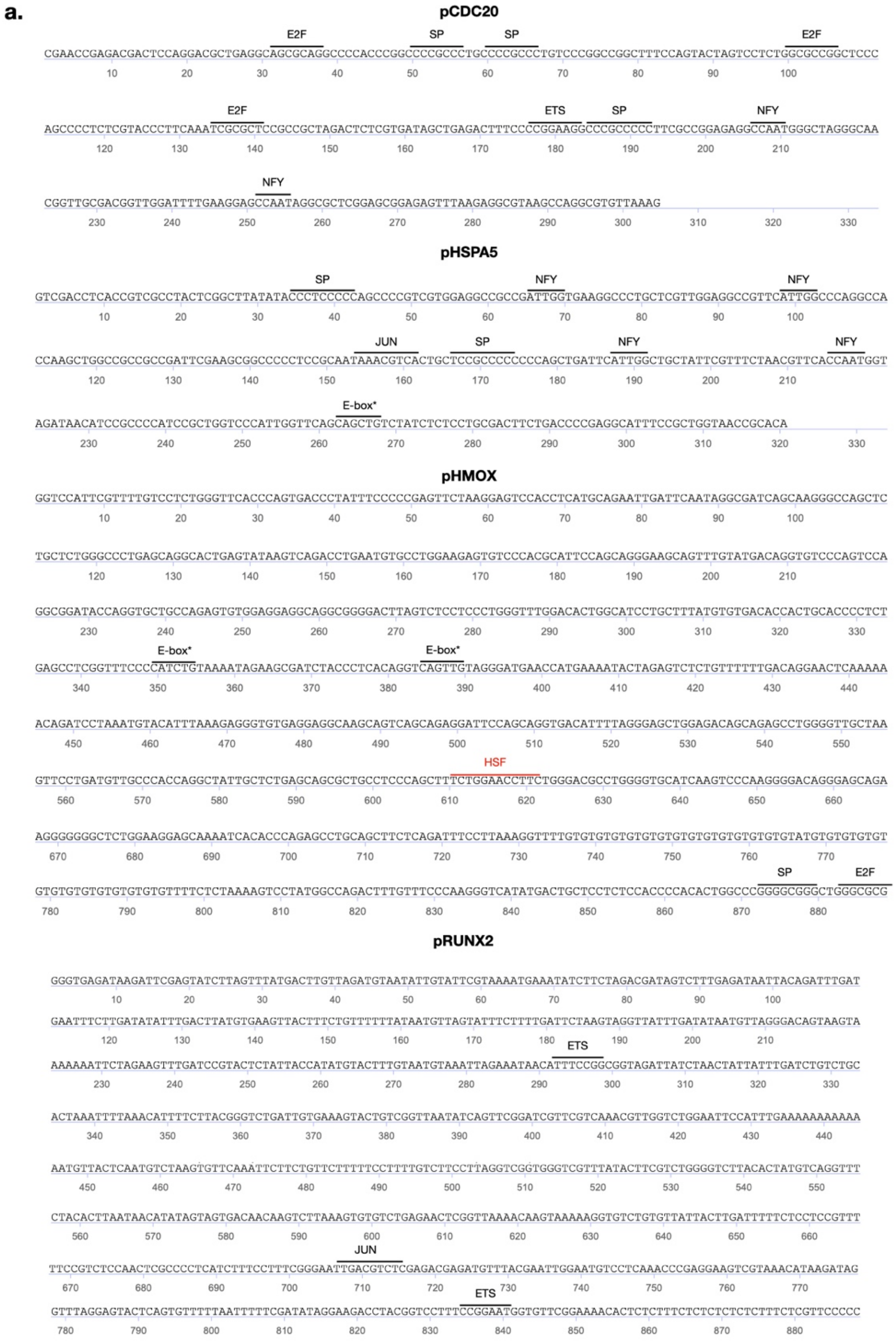

**Supplementary Figure 7. a.** STARR-seq library size distribution after preparation for sequencing. **b.** STARR-seq library complexity quantification for each treatment group. **c.** Landscape of all TFBS in all STARR peaks across different treatment showed that the drug treatment did not alter the global CRE activity to a significant extent. **d.** Normalized motif counts of all EBTF-Co factors after different treatments.

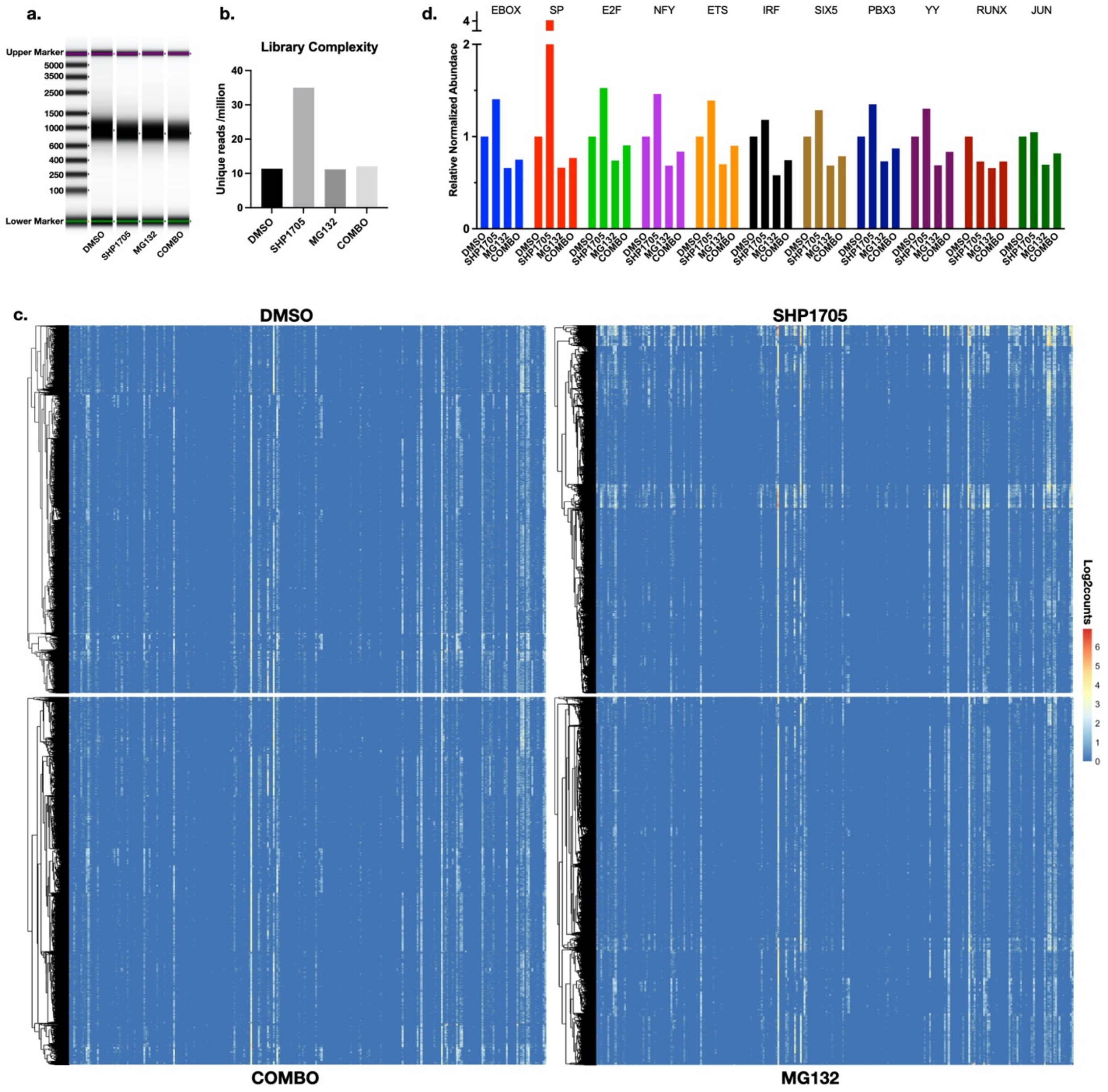

**Supplementary Figure 8. a.** Co-occurrence analysis on 258 non-redundant motif clusters in all treatment groups. **b.** PCA analysis on the motif counting matrix did not reveal meaningful subtypes of the CREs based on their motif counts, and **c.** each PC accounts for no more variance than single dimension. **d.** Contribution of single motifs to the potential repression by drug combination.

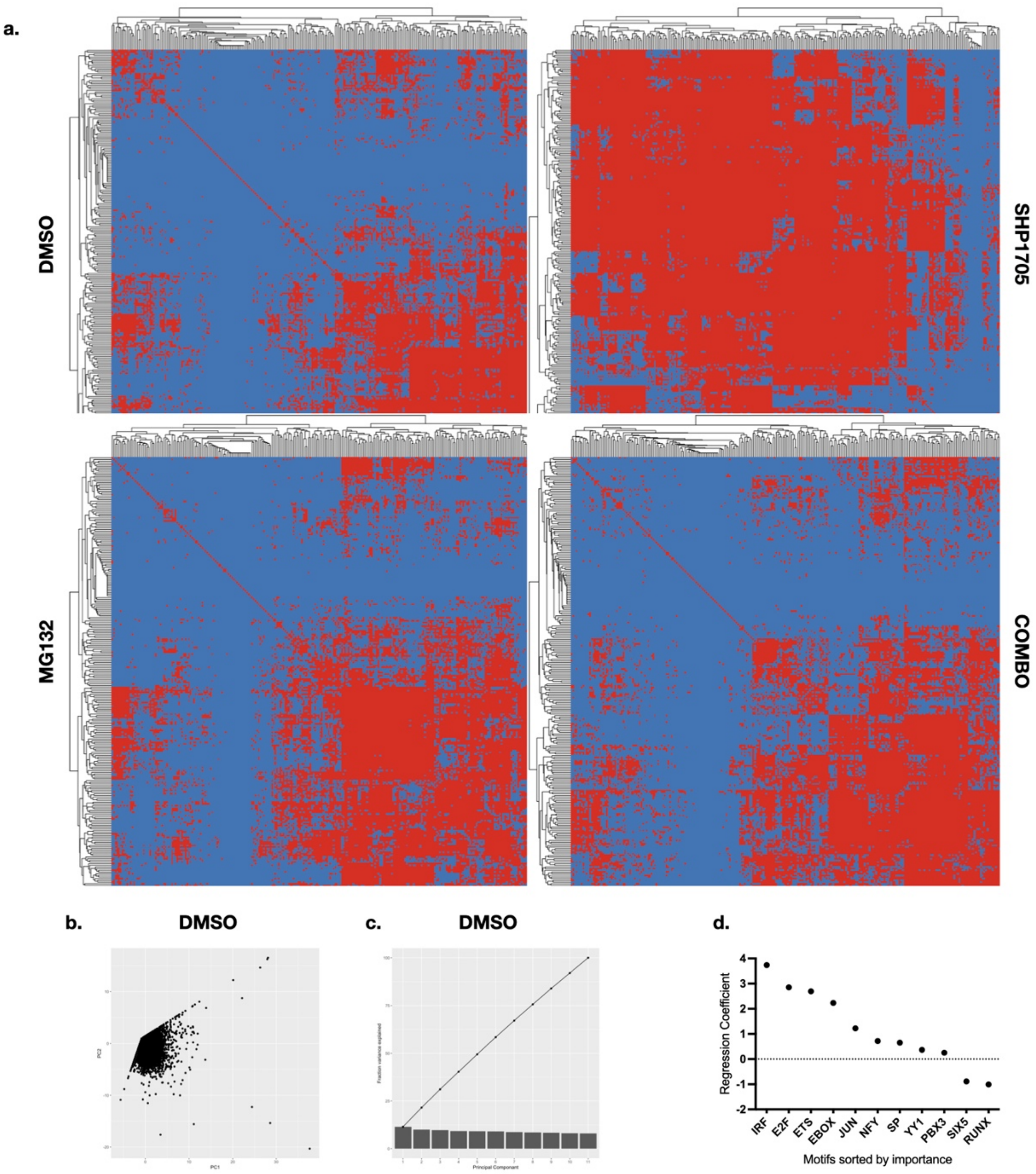

### Supplementary Methods

#### Cell Culture Information

All cell lines were obtained from ATCC. MDA-MB157 (HTB-24), MDA-MB-231 (HTB-26), MDA-MB-436 (HTB-130), MDA-MB-453 (HTB-131), HCC70 (CRL-2315), and HCC 1143 (CRL-2321) were cultured in RPMI 1640 (Invitrogen Cat. 72400120) supplemented with 10% FBS. BT549 (HTB-122) was cultured in RPMI 1640 containing 10% FBS and 0.023 U/mL insulin. Hs-578T was cultured in DMEM with 10% FBS and 0.01 mg/mL insulin.

#### TCGA and METABRIC cohort data analysis

TCGA cohort data was downloaded from UCSC Xena (https://xenabrowser.net/datapages/?cohort=TCGA%20Breast%20Cancer%20(BRCA)&removeHub=https%3A%2F%2Fxena.treehouse.gi.ucsc.edu%3A443). METABRIC data was downloaded from cBioPortal (https://www.cbioportal.org/study/summary?id=brca_metabric).

Differential expression analysis between tumor and normal tissues in TCGA dataset was performed as described in Li et al[1] using `wilcox.test` function. Correlation analysis was done using the R package `corrplot`. Logistic regression was done using the `glm` function in R, and multivariate cox regression was done using the `coxph` function in R with default parameters. Survival analyses were performed using the `survival` and `survminer` packages in R.

#### shRNA lentivirus production

Lentivirus vector clones expressing shBMAL1 and shCLOCK were obtained from Sigma-Aldrich (Sigma). shBMAL1: TRCN0000019097 and TRCN0000019096

(NM_001178.3-1536s1c1, NM_001178.3-689s1c1), shCLOCK: TRCN0000018976 and TRCN0000018978 (NM_004898.2-1053s1c1, NM_004898.2-1494s1c1)

Lentivirus was produced by transfecting the shRNA vector with packaging (psPAX2, Addgene #12260) and envelop (pMD2.G, Addgene #12259) vectors (both are gifts from Didier Trono), into HEK293T cells using Lipofectamine 3000 following manufacturer's protocol. Briefly, HEK 293T cells were seeded in 6-well plate on day 0 to achieve a >90% confluency on the morning of day 1 in 2 mL packaging media (OptiMEM with 5% FBS and Sodium Pyruvate). In the morning of day 1, make Lipofectamine mix (125 µL OptiMEM + 7 µL Lipo 3000 for each well, vortex briefly) and DNA mix (125 µL OptiMEME + P3000 enhancer, then add plasmid DNA) respectively, then mix the two by pipetting to make transfection and incubate at room temperature for 15 minutes. Discard 1 mL of packaging media from each well of HEK293T, then add 250 µL of transfection mix to each well. Six hours after transfection, change the transfection reaction media into fresh packaging media. Twenty-four hours after transfection, the media containing lentivirus was harvested and MOI was determined. Lentivirus were stored at -80 ºC until transduction.

**RNA isolation and Quantitative RT-PCR**

Total RNA of cells was extracted using New England Biolabs Monarch RNeasy Mini kit following manufacturer's manual with on-column DNA digestion. 500 ng of total RNA was used to perform reverse transcription using Super Script IV VILO Master Mix (Invitrogen cat. 11766050) in 10 µL reactions. qPCR was performed using PowerUp SYBR green master mix (Applied Biosystems cat. A25742) in 10 µL reactions (5 µL

Maxter Mix, 3 µL cDNA rxn, 1 µL 5µM fwd +rev primer mix, 1 µL water) in a BioRad thermocycler. All primer sequences are listed in the primer sequence table.

**STARR-seq Library Preparation and Cloning**

STARR-seq library cloning was done as described in the detailed protocol by with minor modifications.[2] hSTARR-seq_ORI vector (Addgene #99296) was obtained from Addgene.

*Genomic Library Insert Generation.* Genomic DNA was extracted from cultured MDA-MB-231 cells using NEB Monarch Genomic DNA Purification Kit (NEB cat. T3010). The genomic DNA (5 µg gDNA in 80 µL water, in 1.7 mL sonification tube) was then sonicated in a Diagenode® BIORUPTOR 300 ultrasonic processor (15 s sonication followed by 15 s pause in 4°C water bath, 3 cycles on high strength), 20 reactions were performed. Fragmented gDNA was run in 1% agarose gel in 20 wells at 140V for 30 minutes. Band of size range 500-750 bp was cut and purified using Macherey-Nagel Gel extraction kit (MN cat. 740609) following manufacturer's protocol. The elution fractions were pooled and cleaned with QIAquick PCR purification kit (QIAGEN cat. 28104), then eluted in 50µL EB buffer. The resulting library was then ligated with Illumina sequencing adaptor using NEBNext Ultra II DNA Library Prep Kit for Illumina (E7645S). End preparation reaction:

| Reaction | | Program | |
|---|---|---|---|
| End Prep Enzyme Mix | 3 µL | 20°C | 30 minutes |
| End Prep Reaction Buffer | 7 µL | 65 °C | 30 minutes |
| Fragmented DNA (500 ng) | 50 µL | 4 °C | Hold |

| | | | |
|---|---|---|---|
| **Total vol.** | 60 µL | Lid | 75 °C |

Ligation reaction:

| **Reaction** | | **Program** | |
|---|---|---|---|
| End Prep Reaction Mixture | 60 µL | 20°C | 15 minutes |
| Ligation Master Mix | 30 µL | 4 °C | Hold |
| Ligation Enhancer | 1 µL | Lid | Off |
| Adaptor for Illumina | 2.5 µL | | |
| **Total vol.** | 93.5 µL | | |

Add 3µL of USER™ Enzyme to the ligation mixture, mix well and incubate at 37 °C for 15 minutes. The reaction was purified twice using Agencourt AMPure XP beads (Beckman Coulter Cat. A63881) following protocols provided by Muerdter and Boryn *el al*.

*Amplification of adaptor-ligated DNA library.* The library insert was amplified by PCR using KAPA HiFi Hotstart ReadyMix (Roche cat. KK2601) in 30 reactions.

| **Reaction** | | **Program** | |
|---|---|---|---|
| Adaptor-ligated DNA library | 1 µL | 98°C | 45 secs |
| Library cloning primer_fwd (10µM) | 2.5 µL | 98 °C | 15 secs |
| Library cloning primer_rev (10µM) | 2.5 µL | 65 °C | 30 secs |
| KAPA HiFi Mix 2X | 25 µL | 72 °C | 45 secs |
| $H_2O$ | To 50 µL | Go to Step 2 | 10 cycles |
| **Total vol.** | 50 µL | 72 °C | 60 secs |

Every 10 PCR reactions were pooled and cleaned with Agencourt AMPure XP beads and QIAquick PCR purification kit following the aforementioned protocol.

*Restriction digest and purification of STARR-seq vector.* 25 µg of STARR-seq screening vector was used for restriction digest in a total of 500 µL reaction (25 µL AgeI-HF, 25 µL SalI-HF, 50 µL CutSmart Buffer (10X), and water up to 500 µL). The reaction was incubated at 37 °C for 2 hours, heat activated at 65 °C for 20 mins, and then run on 1% agarose gel at 140 V for 30 minutes in 10 wells. Th de 3000 bp bands were cut out and extracted using MN Gel and PCR purification kit, then cleaned with QIAquick PCR purification kit and QIAGEN MinElute PCR purification kit (QIAGEN cat. 28006).

*In-fusion HD reaction.* Four reactions were pooled in one tube and a total of 20 reactions were performed.

| **One Reaction** | | **Program** | |
|---|---|---|---|
| Linearized Plasmid | 125 ng | 50°C | 15 minutes |
| PCR amplified library insert | 100 ng | 4 °C | Hold |
| In-Fusion HD Enzyme Premix | 2 µL | | |
| Water | To 10 µL | | |
| **Total vol.** | 10 µL | | |

DNA precipitation was performed for each pooled In-Fusion reaction. The volume of In-Fusion reaction was adjusted to 250 µL using EB buffer, 25 µL 3M NaAc pH5.2 was added and vortexed, then 750 µL ice-cold (-20 °C) 100% ethanol was added, followed by vortex. The mixture was stored at -20 °C for 16 hours. Then DNA precipitation was spin down at full centrifuge speed at 4°C and washed 3 times with 750 µL ice-cold 75% ethanol. After the last wash, the pellets were first dried at room temperature and resuspended in 12.5 µL EB.

*Library expansion.* Cloning reactions are pooled before transformation and distributed to pre-cooled 1.5 mL DNA LoBind tubes (2.5 µL in each tube of total 20 tubes).  20 µL electrocompetent cells (MegaX DH10B$^{TM}$ T1R) to each tube and mixed well with DNA. The mixture was pipetted into pre-cooled cuvettes and electroporated at 2kV, 25 µF, 200 ohms with BioRad GenePulser. Pre-warmed recovery media (1mL per cuvette) was added immediately after electroporation, and cells were recovered at 37 °C for 1 hour. All transformation reactions were pooled after recovery and distributed evenly to 12 L pre-warmed LB media with 100 µg/mL Ampicillin. The cells were cultured overnight while shaking at 37 °C till the OD600 is between 2-2.6. Bacteria cells were spun down and pooled, DNA library was extracted using QIAGEN Plasmid Giga Kit (QIAGEN cat. 12991) following manufacturer's protocol.

**STARR-seq Screening**

MDA-MB-231 cells were cultured in 15 cm tissue culture plates and reached 90% confluency on the day of transformation (5 plates, total 1 x $10^8$ cells for each treatment group). Before transformation, drugs corresponding to each group (0.1% DMSO, 10µM SHP1705, 100 nM MG132, and combination) were diluted in fresh media without FBS or antibiotics and exchanged the culture media. Transformation was done with Lipofectamine 3000 following manufacturer's protocol. For each 15 cm plate, 2.5 µg STARR library DNA was diluted into 1250 µL OptiMEM, then 50 µL P3000 enhancer was added to make the DNA mix. Then 1250 µL OptiMEM containing 65 µL well mixed Lipofectamine 3000 was add to the DNA mix and mixed by pipetting. The transformation reaction was incubated at room temperature for 15 minutes and added dropwise into the cells containing drugs. Eight hours after transformation, cells were harvested using TrypLE treatment, and total RNA was isolated with QIAGEN RNeasy Maxi Kit (QIAGEN cat. 75162) following manufacturer's protocol.

mRNA was isolated from total RNA using Oligo-dT magnetic beads (Dynabeads Oligo(dT)$_{25}$, Invitrogen cat. 61005) following manufacturer's protocol. Briefly, the beads were washed first with Binding Buffer and resuspended to the volume of the RNA solution to be purified. Then equal volume of RNA was added and incubated on a rolling shaker for 10 minutes at room temperature. The samples were then put on magnet for 2 minutes at room temperature, and the supernatants were discarded. The beads were washed with Washing Buffer B, then RNA was eluted in 10 mM Tris-HCl on an 80 °C heating block for 3 minutes shaken at 750 rpm. Selected RNA was cleaned using NEB Monarch RNA miniprep kit with on-column DNA digestion.

Reverse transcription (RT) was done with SuperScript III (Invitrogen cat. 18080093). For each treatment group, 5 µg PolyA+ RNA was used to perform RT in 2 PCR tubes (2.5 µg each tube). The reactions are set up per the following table:

| **RNA Mix** | | **RT Mix** | |
|---|---|---|---|
| polyA$^+$ RNA | 2.5 µg | 10X RT buffer | 10 µL |
| GSP (2 µM) | 5 µL | 25 mM $MgCl_2$ | 20 µL |
| dNTP (10 mM) | 5 µL | 0.1 M DTT | 10 µL |
| Water | To 50 µL | RNase OUT | 5 µL |
| | | SuperScript III RT | 5 µL |
| **Total vol.** | 50 µL | **Total vol.** | 50 µL |

RNA mixes were prepared first, and RT mixes were added to make the reaction. Program: 50 °C for 1 hour, 85 °C for 5 minutes, hold at 4 °C. 1 µL RNase H was added to each tube and incubated at 37 °C for 1 hour. RT reactions were purified using AMPure XP beads.

Sequencing library was prepared in two steps. First, four junction PCR (jPCR) reactions were performed for each treatment group.

| **One jPCR Reaction** | | **Program** | |
|---|---|---|---|
| cDNA | 20 µL | 98 °C | 45 s |
| Primer (j-fwd + j-rev, 5 µM each) | 5 µL | 98 °C | 15 s |
| KAPA HiFi 2X | 25 µL | 65 °C | 30 s |
| | | 72 °C | 45 s |
| | | Go to Step 2 | 14 times |
| **Total vol.** | 50 µL | 72 °C | 60 s |

jPCR reactions were purified with AMPure beads and eluted in nuclease-free water. Then five sequencing-ready PCR reactions were performed for each treatment group. Index primers are from NEBNext Multiplex Oligos for Illumina (Index Primers Set 1, NEB cat. E7335). Index Primer 2,4,6, and 12 were used for treatment group DMSO, SHP1705, MG132, and COMBO respectively.

| **One seq-ready PCR Reaction** | | **Program** | |
|---|---|---|---|
| DNA from cleaned jPCR | 20 µL | 98 °C | 45 s |
| Universal PCR Primer (µM) | 2.5 µL | 98 °C | 15 s |
| Index Primer (10 µM) | 2.5 µL | 65 °C | 30 s |
| KAPA HiFi 2X | 25 | 72 °C | 45 s |
| | | Go to Step 2 | 9 times |
| **Total vol.** | 50 µL | 72 °C | 60 s |

Sequencing-ready library was purified using SPRI beads (Beckman cat. B23318).

#### STARR-seq data analysis

Library complexities were calculated using the `preseq` program (v.2.0.0).[3] After peak calling, motif counts were done with the `countPWM` function in the `Biostrings` package in R with threshold set to 80%. Normalized abundance of single motifs was calculated by the following formula:

$$Normalized\ Abundance = \frac{number\ of\ motifs * peak\ pileup}{number\ of\ peaks * complexity\ of\ library}$$

and normalized to DMSO group to obtain relative normalized abundance. Logistic regression was done using the `glm` function in R.

### References


1. Li, Y., Ge, X., Peng, F., Li, W. & Li, J. J. Exaggerated false positives by popular differential expression methods when analyzing human population samples. *Genome Biol* **23**, (2022).
2. Muerdter, F. *et al.* Resolving systematic errors in widely used enhancer activity assays in human cells. *Nat Methods* **15**, (2018).
3. Daley, T. & Smith, A. D. Predicting the molecular complexity of sequencing libraries. *Nat Methods* **10**, (2013).

| qPCR Primers | | |
|---|---|---|
| | Forward | Reverse |
| PPIA | CAACCCCACCGTGTTCTTCG | GTGAAGTCACCACCCTGACAC |
| DBP | TGACCCTCGAAGACATCGCT | TCGTTGTTCTTGTACCGCCG |
| CRY1 | CTTCCTGACACGAGGGGACC | GCCACATCCAACTTCCAGCA |
| PER1 | TCCATTCGGGTTACGAAGCT | GCAGCCCTTTCATCCACATC |
| STARR-seq library preparation primers | | |
| in-fusion forward | TAGAGCATGCACCGGACACTCTTTCCCTACACGACGCTCTTCCGATCT | |
| in-fusion reverse | GGCCGAATTCGTCGAGTGACTGGAGTTCAGACGTGTGCTCTTCCGATCT | |
| RT Gene-specific primer | CTCATCAATGTATCTTATCATGTCTG | |
| junction PCR forward | TCGTGAGGCACTGGGCAG*G*T*G*T*C | |
| junction PCR reverse | CTTATCATGTCTGCTCGA*A*G*C | |
